# Endogenous retroviruses co-opted as divergently transcribed regulatory elements shape the regulatory landscape of embryonic stem cells

**DOI:** 10.1101/2021.06.11.448013

**Authors:** Stylianos Bakoulis, Robert Krautz, Nicolas Alcaraz, Marco Salvatore, Robin Andersson

## Abstract

Transposable elements are an abundant source of transcription factor binding sites and favorable genomic integration may lead to their recruitment by the host genome for gene regulatory functions. However, it is unclear how frequent co-option of transposable elements as regulatory elements is, to which regulatory programs they contribute and how they compare to regulatory elements devoid of transposable elements. Here, we report a transcription initiation-centric, in-depth characterization of the transposon-derived regulatory landscape of mouse embryonic stem cells. We demonstrate that a substantial number of transposable elements, in particular endogenous retroviral elements, carry open chromatin regions that are divergently transcribed into unstable RNAs in a cell-type specific manner, and that these elements contribute to a sizable proportion of active enhancers and gene promoters. We further show that transposon subfamilies contribute differently and distinctly to the pluripotency regulatory program through their repertoires of transcription factor binding sites, shedding light on the formation of regulatory programs and the origins of regulatory elements.

## Introduction

Transcriptional regulatory elements are stretches of genomic sequence that exert enhancer and promoter activities essential for the precise spatial and temporal control of gene expression (Haberle and Stark 2018; Beagrie and Pombo 2016; Shlyueva et al. 2014; Andersson and Sandelin 2020). The activity of a regulatory element is controlled by the specificity of transcription factors (TFs) to bind the element and the density of their binding sites, both of which are in turn dependent on its DNA sequence (Nguyen et al. 2016; Weingarten-Gabbay et al. 2019; Smith et al. 2013; Grossman et al. 2017). TF DNA-sequence preferences (Nitta et al. 2015), gene expression (Chan et al. 2009; Berthelot et al. 2018), and the specific regulation of genes by TFs (Odom et al. 2007; Schmidt et al. 2010) are generally well conserved across eukaryotes. On the contrary, regulatory elements with enhancer activity are generally associated with high evolutionary turnover (Vierstra et al. 2014; Young et al. 2015; Villar et al. 2015).

Transposable elements (TEs) are an abundant source of TF binding sites that contribute to the spread of sequences with regulatory potential (Chuong et al. 2017). In mammals, the large majority of TEs are dormant, having lost their ability to replicate and are maintained repressed by H3K9me3, H3K27me3, and DNA methylation (Walter et al. 2016; Maksakova et al. 2013; Matsui et al. 2010; Karimi et al. 2011; Rowe et al. 2013). However, favorable genomic integration may lead to the co-option of TEs as endogenous regulatory elements by utilizing their native or acquired TF binding sites (Bourque et al. 2008; Chuong et al. 2013; Sundaram et al. 2014; Trizzino et al. 2017; Sundaram et al. 2017; Sun et al. 2018; Barakat et al. 2018; Cao et al. 2019; Todd et al. 2019; Miao et al. 2020). Consequently, the majority of species-specific open chromatin regions are associated with TEs (Vierstra et al. 2014) and TE-derived enhancers are generally not well conserved across evolution (Jacques et al. 2013; Sundaram et al. 2014; Glinsky and Barakat 2019). In embryonic stem cells (ESCs) long terminal repeat (LTR) retrotransposons bound by pluripotency TFs, including OCT4 and NANOG, have been shown to possess enhancer activities (Barakat et al. 2018; Todd et al. 2019) and initiate transcription (Fort et al. 2014). In addition, species-specific TE-derived regulatory elements cause binding differences of pluripotency TFs between human and mouse ESCs (Kunarso et al. 2010). Distribution and fixation of mobile elements with readily available regulatory potential may therefore provide regulatory innovation but also stabilize the gene regulatory functions of TFs. Thus, characterizing the contribution of TEs to transcriptional regulation has the potential to provide insights into the formation of pluripotency regulatory programs, the origins of regulatory elements and thus the basis for their evolutionary turnover.

Most studies to date have inferred TE-associated regulatory elements from chromatin-accessible loci flanking nucleosomes with specific histone modifications indicative of enhancers (e.g., H3K4me1, H3K27ac). However, only few of such predicted loci show enhancer activity (Kheradpour et al. 2013; Todd et al. 2019). Rather, a growing body of literature points toward divergent transcription initiation as a key property of active regulatory elements with either enhancer or promoter function (Kim et al. 2010; Andersson et al. 2014b, 2014a, 2015a; Scruggs et al. 2015; Chen et al. 2016; Rennie et al. 2018; Henriques et al. 2018; Andersson and Sandelin 2020). While there is a general relationship between histone modifications and the transcriptional output of a regulatory element (Andersson and Sandelin 2020; Core et al. 2014; Henriques et al. 2018; Rennie et al. 2018), the transcriptional status of a regulatory element better reveals its regulatory potential (Andersson et al. 2014a; Wu et al. 2014; Rennie et al. 2018). Characterization of TE-derived enhancers from transcription initiation events therefore provide a more accurate picture of their regulatory contribution.

Divergent transcription of regulatory elements is established at closely spaced pairs of divergently oriented core promoters within open chromatin, resulting in long non-coding enhancer RNA (eRNA) transcripts at regulatory elements with enhancer activity, and pairs of mRNAs and promoter upstream transcripts (PROMPTs) at gene promoters (Koch et al. 2011; Andersson et al. 2014b, 2014a; Scruggs et al. 2015; Andersson et al. 2015a; Rennie et al. 2018). Such transcription initiation events are accurately identified and quantified through 5’ end sequencing of capped RNAs (CAGE, Cap Analysis of Gene Expression) (Takahashi et al. 2012; Kawaji et al. 2014). eRNAs and PROMPTs are generally non-polyadenylated and thus unprotected at their 3’ ends, and as a consequence these RNA species are frequently targeted by the 3’-5’ ribonucleolytic RNA exosome for degradation (Preker et al. 2008; Ntini et al. 2013; Andersson et al. 2014b).

Although co-option of TEs as regulatory elements has been established as a mode of regulatory innovation, its extent and how TE-derived regulatory elements compare to non-TE associated regulatory elements (e.g., with regards to divergent transcription initiation, RNA metabolism and TF binding) remain unclear. Encouraged by the possibility to study TE-associated regulatory elements through transcription initiation mapping (Fort et al. 2014), we here investigate how the wide repertoire of mouse TEs contribute to the transcription initiation landscape and thus regulatory elements of mouse ESCs (mESCs). We demonstrate that many dormant TE insertions, in particular endogenous retroviral elements (ERVs), carry open chromatin regions that are divergently transcribed into unstable RNAs targeted by the exosome for degradation. Furthermore, open chromatin regions that are associated with transcribed ERVs show a high degree of species specificity and a large fraction of these either have enhancer function or contribute to gene promoters. This suggests that TE co-option as regulatory elements contributes to a sizable proportion of active species-specific regulatory elements in mESCs. Our transcription-centric approach allows for an unbiased systematic investigation of the regulatory potential across TE subfamilies, indicating that these contribute differently to the TF binding repertoire of the mouse genome, which can be linked to regulatory specificity in mESCs.

## Materials and methods

### E14 mESC and HeLa S2 CAGE libraries, data processing and mapping

Previously sequenced E14 mESCs, samples after 3 days of differentiation of mESCs into embryoid bodies (GEO ID GSE115710) (Lloret-Llinares et al. 2018) and human HeLa S2 cells (GEO ID GSE62047) (Andersson et al. 2014b) CAGE (Takahashi et al. 2012) libraries were collected. As described in each report, CAGE libraries were prepared from exosome depleted samples as well as from control samples. E14 mESCs and differentiated embryoid bodies were transduced with pLKO vectors encoding the shRNA: SHC002 (scrambled control -referred to as Scr control) and <u>NM_025513</u>.1-909s1c1 (referred to as Rrp40 exosome knockdown). HeLa cells were transfected with enhanced green fluorescent protein (eGFP - referred to as EGFP control) and the hRRP40 (*EXOSC3*) siRNA (referred to as RRP40 exosome knockdown). The CAGE libraries from mouse and human were processed as in the original publications, with some minor modifications. Reads were trimmed using the FASTX-Toolkit (Version 0.013 - http://hannonlab.cshl.edu/fastx_toolkit/) to remove linker sequences (Illumina adaptors) and then filtered for a minimum sequencing quality of 30 in 50% of the bases. Mapping to the mouse reference genome (mm10) was performed using Bowtie (Version 1.1.2), applying the following parameters to ensure several (up to 100) good alignments per read, which is essential for the rescue and analysis of TE-derived sequences: *-k 100* (report up to 100 good alignments per read), *-m 100* (eliminate reads that map > 100 times), *--best* and *--strata* (report alignments which have the highest quality). Reads that mapped to unplaced chromosome patches or chrM were discarded. Finally, all reads corresponding to reference rRNA sequences (mouse: <u>BK000964.3</u>, human: <u>U13369.1</u>) with up to 2 mismatches were discarded using rRNAdust (<u>fantom.gsc.riken.jp/5/suppl/rRNAdust/</u>). Mapping to the human reference genome (hg19) was performed using the exact same parameters and version of Bowtie.

For exploratory/comparative purposes, the mapping of HeLa S2 CAGE libraries was performed with 3 additional alignment approaches (Supplementary Fig. 1 B,C). BWA (Burrows-Wheeler Aligner) version 0.7.15-r1140 (Li and Durbin 2009) was used with the parameter -n 2 (maximum distance for each alignment) and a subsequent mapping quality threshold (MAPQ >20), using SAMtools v1.3.1 (Li et al. 2009). BWA-PSSM, a modification of BWA using position specific scoring matrices (PSSM) (Kerpedjiev et al. 2014), was used with parameters -n 2, -m 2000 (to allow for more suboptimal partial alignments to be tested) and a downstream MAPQ>20 threshold. Finally, LAST version 801 (Kiełbasa et al. 2011) was used with parameters -u NEAR -R 01 (*lastdb*) and -Q 1 and -D 100 (*lastal*), followed by a downstream MAPQ>20 threshold.

### Probabilistic multi-mapping rescue of CAGE tags

Following initial mapping of reads with Bowtie, we employed *MUMRescueLite* (Hashimoto et al. 2009) to resolve short multi-mapping CAGE reads that aligned equally well to more than one genomic location. In short, this method examines the information about the local context of potential mapping positions given by uniquely mapping reads. By assuming that multi-mapping reads are more likely to come from regions which already have more uniquely mapping reads, MUMRescueLite probabilistically assigns the true source of a multimapping read. The probabilistic scheme behind the tool is weighted by the abundance and location of uniquely mapping reads. Thus, a nominal window parameter is required for which to identify unique mappers that occur around (upstream and downstream) each locus occupied by a multi mapper. In addition, a weight threshold is required over which one locus of a multi mapper is “rescued”, referring to the fraction out of the total number of unique mappers proximal to all loci associated with a specific multi mapper. The window parameter was cautiously selected to be 50 bp after a saturation investigation of how many reads are rescued. The weight threshold was set to 0.7, in order to select one locus as the true source of a multi-mapping read.

### TElocal-inferred expression of mESC TE families

For statistical comparison purposes, CAGE unique and multi-mapping reads aligned with Bowtie, as described above, were supplied to TElocal v 0.1.0 from the TEToolkit suite (Jin et al. 2015) in order to quantify transposable element expression at the locus level. The resulting quantification for each TE for both TElocal and multi mapping rescuing output was normalized to tags per million mapped reads (TPM) and the results of the two approaches were compared using Spearman’s rank-based correlations.

### CAGE tag clustering, quantification, and normalization

Following multi-mapping rescue, the number of overall CAGE tag 5′ends were counted for each genomic position to obtain base-pair (bp) resolution of CAGE transcription start sites (CTSSs). We then assigned CTSSs to transposable elements (TEs) as defined by RepeatMasker v4.0.7 (http://www.repeatmasker.org/) and considered only instances with two or more CAGE tags. TEs denoted as Alu elements in RepeatMasker were considered B1 and proto-B1 (PB1) elements. Tag clusters (TCs) were generated from pooled CAGE libraries per condition, and wide TCs were narrowed or decomposed into sub-peak TCs if containing multiple peaks (https://github.com/anderssonlab/CAGEfightR_extensions), as previously described (Rennie et al. 2018). In short, CTSSs located within 20 bp from each other on the same strand were merged into initial TCs. For each TC, the bp with the most abundant count (summit position) or the median of multiple equal summits were identified. Next, the fraction of total CTSSs of each location within a TC to that of the summit position was calculated. If a position carried less than 10% of the summit signal, it was discarded and TCs split were merged if positioned within 20bp from each other on the same strand. Expression quantification of each individual TC in each CAGE replicate took place by adding up the CTSSs falling into them. Using the CAGE genomic background noise estimation (as described below), all TCs with expression values below the noise threshold were discarded from further analyses. Expression levels of TCs were normalized to tags per million mapped reads (TPM). Finally, we assigned normalized TCs to TEs on the same strand through direct overlap using BEDtools (Quinlan and Hall 2010).

### Footprints of CAGE expression on TEs

To investigate the relationship between transposable element families and/or classes to CAGE TSS locations, we plotted the average binarized (presence or absence of CTSS, regardless of expression value) pooled CTSS signal 500bp upstream, across the body and 500bp downstream of each TE instance using deepTools (Ramírez et al. 2016). Unique TE instance profiles were averaged for each TE family or class based on their Repeatmasker annotation. Furthermore, we constructed a synthetic CAGE uniqueness track by mapping the mm10 reference genome split in 25 bp long segments back to itself. Localization of the synthetic CAGE tags on a bp resolution was conducted as described above at the CTSS level, assigned to TE instances and the signal was binarized, representing the expected uniquely mapped background signal. The log_2_ ratio of observed (CAGE libraries) versus expected (as estimated from the synthetic uniqueness track) average binarized signal was calculated in R (http://www.R-project.org/).

### HeLa RNA-seq data processing

RNA-seq data from HeLa cells depleted of hRRP40 using siRNA-mediated knockdown as described elsewhere (Andersen et al. 2013) (SRA accession: SRX365673) were considered. Briefly, after filtering of low-quality reads, removal of Illumina adaptors and reads shorter than 25 bp with Trimmomatic v0.36 (Bolger et al. 2014), reads were mapped against the human reference genome (hg19) using HISAT2 v2.1.0 (Kim et al. 2015). Uniquely mapped and properly paired reads were selected with SAMtools v1.3.1. Gene-level expression quantification of mapped reads was performed with featureCounts v1.6.3 (Liao et al. 2014). Further analyses and comparison to gene-level expression with CAGE using generalized linear Poisson regression models with backward elimination for variable selection was performed in R using the glm() function (see Supplementary note).

### CAGE gene-level expression quantification

For statistical comparison of gene-level HeLa RNA-seq expression to gene-level HeLa expression as measured by CAGE, we quantified abundances of genes using CTSSs within +/- 500 bp windows from the 5’end of GENCODE primary mouse sequence and annotation (GRCm38) version M10 transcripts (Frankish et al. 2019), as CAGE signal from HeliScopeCAGE saturates after ∼500bp from annotated gene TSSs (Kawaji et al. 2014). Gene-level abundances were quantified by first merging potentially overlapping TSS-centered windows per transcript belonging to the same gene and then summing the expression levels of all transcript windows for each gene. Similarly, gene-level expression was quantified using CAGE data for mESC and embryoid bodies.

### Processing of DNase-seq data and DHSs as focus points for transcription initiation

For identification of TE-associated regulatory elements, sequencing reads from DNase-seq for the mouse ES-E14 cell line (GEO ID GSE37073 / GSM1014154) were processed using the ENCODE DNase-HS pipeline. Called hotspot FDR 1% peaks in the mouse reference genome (mm10) were used as DNase I hypersensitive sites (DHSs). DHSs were used as focus points of minus and plus strand expression by defining DHS midpoints as positions optimizing the coverage of proximal CAGE tags within flanking windows of size +/- 300 bps around them, as previously described (Rennie et al. 2018). The final set of 165,052 transcribed DHS was determined by filtering DHSs to not overlap any other DHS +/- the 300bp window on the same strand and to be supported by either control or exosome knockdown CAGE expression above the noise threshold (described below). Transcriptional directionality and exosome sensitivity scores were calculated considering this set of DHS regions, as defined previously (Andersson et al. 2014b). In short, the directionality score measures the expression level strand bias within transcribed DHSs, it ranges from -1 to 1 (100% minus or plus strand expression), while 0 indicates balanced bidirectional transcription. The exosome sensitivity score measures the relative amount of degraded RNAs by the exosome by quantifying the fraction of exosome-depleted CAGE expression seen only after exosome depletion: exosome sensitivity values closer to 1 are indicative of highly unstable RNAs.

### Estimation of CAGE genomic background noise

The estimation of a CAGE genomic background noise threshold (https://github.com/anderssonlab/CAGEfightR_extensions) for robust assessment of lowly expressed regions was based on quantifying the CAGE 5’ends in randomly selected uniquely mappable regions of 200 bp distal to known TSSs, exons and DHSs, followed by extracting the 99th percentile of the empirical distribution of CAGE expression and using the max value across control libraries as a noise threshold for significant expression in further analyses, described in detail elsewhere (Rennie et al. 2018).

### Genomic annotation of transcribed TEs

We annotated TE-associated TCs based on different genomic regions as defined using GENCODE version M10 and BEDtools, ensuring there are no overlapping regions counted twice. Coordinates for all genic regions (exons, 5’ UTRs, 3’UTRs) were extracted from the GENCODE annotation. Promoter regions were defined as regions at the starting positions of each transcript +/- 500 bp. To define intronic regions, we subtracted the exonic regions from the genic regions. Finally, distal/intergenic regions were defined as the remaining parts of the genome in-between annotated genes.

### Estimation of evolutionary conservation of TE-associated DHSs

Genomic regions spanning +/- 150 bp around mESCs DHS signal peaks carrying CAGE tags and overlapping TEs were aligned to rat (rn7) and human (hg38) assemblies using the *UCSC liftOver* tool (Hinrichs et al. 2006) with a -minMatch=0.6 parameter. Similarly, 300bp regions of nonTE-associated DHSs carrying CAGE tags, the mm10 genome assembly split in 300bp fragments, the subset of those fragments not overlapping TEs, and all TE instances of RepeatMasker (full length) were aligned to rat and human assemblies. The genomic regions that had a >60% match (coverage) with those in the other species were considered orthologous.

### FANTOM enhancers and enhancer peak calling from STARR-seq data

Transcribed enhancers identified by remapped FANTOM5 CAGE libraries to mm10, deposited in Zenodo (Dalby et al. 2018), were associated with TE-associated DHSs by overlap using BEDTools. STARR-seq data for 2iL grown mESCs E14Tg2a (E14) (Peng et al. 2020) (GEO ID GSE143546) were used to evaluate the enhancer potential of transcribed TEs. STARR-seq data were processed using Bowtie2 (Langmead and Salzberg 2012) and SAMtools (Li et al. 2009). Reads were aligned to the mouse reference genome (mm10) using Bowtie2 (*--very sensitive*). The reads of the two replicates from each sample were sorted and merged and reads falling into regions from the ENCODE blacklist of the mouse reference genome were removed. STARRPeaker (Lee et al. 2020) was used to identify potential enhancers with default parameters and an adjusted p-value threshold of 0.05. Potential enhancers called from STARR-seq data were associated with expressed TE-associated DHSs by overlap using BEDTools.

### Processing and analysis of histone modification ChIP-seq data

Mouse ENCODE E14Tg2a or E14 mESCs ChIP-seq data for six histone modifications: H3K4me3, H3K4me1, H3K27ac, H3K36me3, H3K9me3 (GEO ID GSE136479) and H3K27ac (GEO ID GSE31039) were processed using the ENCODE ChIP-seq processing pipeline (version 1.3.6). Adapters and low quality reads were filtered with cutadapt v2.5 (Martin 2011), reads were mapped to the mouse genome assembly mm10 with bwa, duplicate reads were removed with Picard v2.20.7 (http://broadinstitute.github.io/picard/), ENCODE mm10 blacklist regions were masked and only reads with mapping quality above 30 were considered for further downstream analyses. For the heatmap and footprint plots, ChIP signal expressed as fold-over input control was averaged across sites in 10 bp bin intervals from the CAGE TC summit position up to a maximum of +/- 2000bp, using deepTools version 3.1.3. Hierarchical clustering, annotation and visualization were conducted in R with ChIPSeeker (Yu et al. 2015), *ggplot2* (https://ggplot2.tidyverse.org) and *profileplyr* (R package version 1.6.0), using clustering parameters *“rowMax’’* for summarizing the ranges across samples and *“median”* for defining proximity between clusters. The association of clusters to TE classes and families was done in R, using the Fisher’s exact test with Bonferroni correction for multiple testing.

### Chromatin state discovery and characterization using chromHMM

To annotate TE-associated DHSs and characterize which regulatory elements in mESCs (mm10) they are occupying, we constructed a 12-state model chromatin states map to identify genomic regions enriched in specific combinations of histone modifications and TF marks, as previously described (Pintacuda et al. 2017). The multivariate hidden Markov model framework of chromHMM (Ernst and Kellis 2012, 2017) was applied to the mouse reference genome (mm10) and ENCODE ChIP-seq data in E14 cells for the following ten marks: H3K4me1, H3K4me3, H3K27me3, H3K27ac, H3K9me3, H3K9ac, H3K36me3, CTCF, Nanog, Oct4. TE-associated DHSs and non-TE DHSs associated with TCs from control and exosome-depleted CAGE libraries were overlapped with the chromHMM states with BEDtools and the heatmaps were generated in R using heatmap.2 from the *gplots* package (R package version 3.1.1.).

### Transcription factor motif analysis

We scanned full ERV transposons carrying > 1 CAGE tag with known TF motifs in the HOMER motif database using findMotifsGenome.pl (Heinz et al. 2010). For each TF, findMotifsGenome.pl employs a hypergeometric test to compare the number of motifs found in the target set with that found in a specified background set. The tool was run using the set of expressed ERV transposons per subfamily as the target set and the full set of non-TE-associated CAGE TCs (+/- 200bp regions) as the background set. The significantly enriched TFBS motifs were selected with three additional conditions: (1) TF genes are expressed in mESCs using gene-level CAGE quantification; (2) at least 10% of the target set contained the motif; and (3) known TF genes have a match score > 0.9 to the *de novo* motifs found by HOMER. The motif enrichment score was calculated as log_2_ (% of target sequences with motif / % of background sequences with motif). Heatmaps of TF motif enrichment were generated in R using the *ggplot2* package. In order to account for cell type specific TE expression, as measured by CAGE in mESCs and embryoid bodies, we scanned TE-associated DHSs in mESCs, using scanMotifGenomeWide.pl (Heinz et al. 2010), with the position weight matrices (PWMs) of nine pluripotency TFs that demonstrated enrichment across several transcribed ERV subfamilies (Fig 5A). ERV instances carrying a predicted binding site of at least one out of the nine TFs were considered to be associated with pluripotency TFs. Comparisons between binding sites of selected mouse TFs were performed using TomTom of the MEME suite including the three mouse-specific motif databases: HOCOMOCO Mouse (v11 FULL), UniPROBE Mouse (Sci09 Cell08), Embryonic Stem Cell TFs (Chen2008) both as query and target motifs.

### Functional Enrichment Analysis

Gene ontology (GO) and pathway enrichment analyses, using TE-regulated ABC (activity-by-contact) predicted target genes (retrieved from https://osf.io/uhnb4/) for enhancer elements (Fulco et al. 2019) as target sets and all ABC predicted target genes as background, were performed in R using the package *clusterProfileR* (Yu et al. 2012). Only statistically significant results (p.adjust < 0.05) were considered after employing false discovery rate for multiple testing correction. ABC region coordinates were lifted from mm9 to mm10 using the UCSC liftOver tool (Rhead et al. 2010) and were associated with CAGE TCs by coordinate overlap (BedTools).

## Results

### Transcription at transposable elements is divergent and results in unstable RNAs

To investigate the prevalence and co-option of TEs as regulatory elements, we first characterized the association of TEs with divergent transcription initiation, a key property of regulatory elements with either enhancer or promoter activities (Andersson et al. 2014a, 2015b; Andersson and Sandelin 2020; Rennie et al. 2018; Mikhaylichenko et al. 2018; Henriques et al. 2018). To this end, we analyzed the genomic location of sequencing reads of capped RNA 5’ ends in mESCs. 5’ ends of capped RNAs reveal transcription start sites (TSSs) of RNA polymerase II transcripts and can be accurately assayed using CAGE (Takahashi et al. 2012). Here, we considered CAGE data from mESCs depleted for the RNA exosome core component Rrp40 (Lloret-Llinares et al. 2018). TSSs identified by this data thus also include those whose RNAs are targeted by the exosome for degradation.

To allow characterization of TSSs at base-pair resolution within TE insertions, we considered both uniquely mapping and multi-mapping rescued (Faulkner et al. 2008; Hashimoto et al. 2009) CAGE reads (see Methods). Multi-mapping rescue increased the number of identified TE-associated TSSs and improved expression quantification of TE-associated loci in mESCs and, as a confirmation, also in HeLa cells (Supplementary note; Supplementary Figs. 1-4). Comparable expression levels were observed using an alternative strategy based on maximum likelihood alignments to annotated repeats (Jin et al. 2015), although the sets of detected, expressed TE insertions varied between methods (Supplementary note; Supplementary Fig. 5).

The ability to infer the TSSs and expression levels of TE-derived RNAs from CAGE prompted us to characterize the transcription initiation patterns across TE families. 82,383 mouse TE insertion events were associated with measurable RNA polymerase II transcripts in mESCs. Among these, endogenous retroviral TE family elements (ERV1, ERVK, ERVL, ERVL-MaLR) were most prevalent (p<2e-16, Fisher’s exact test; Supplementary Fig. 2A). These TE families and L1 LINE elements showed preferential TSS locations close to their repeat body boundaries (Fig. 1A,B). Of note, TSS location preferences could in general not be explained by varying mappability over repeat bodies (Supplementary Fig. 6). However, a lack of detected expression in DNA transposons could be due to low mappability, suggesting that expression quantification and TSS mapping of this class may require alternative approaches or longer sequencing reads. By taking mappability into account, a strong divergent pattern for L1 elements originating at their 5’ and 3’ ends was revealed (Supplementary Fig. 6), suggesting that some of these may utilize their native TSSs. We note, however, that TSSs within TEs often deviate from their native TSSs, as seen for ERVs (Supplementary Fig. 6).

**Figure 1:**
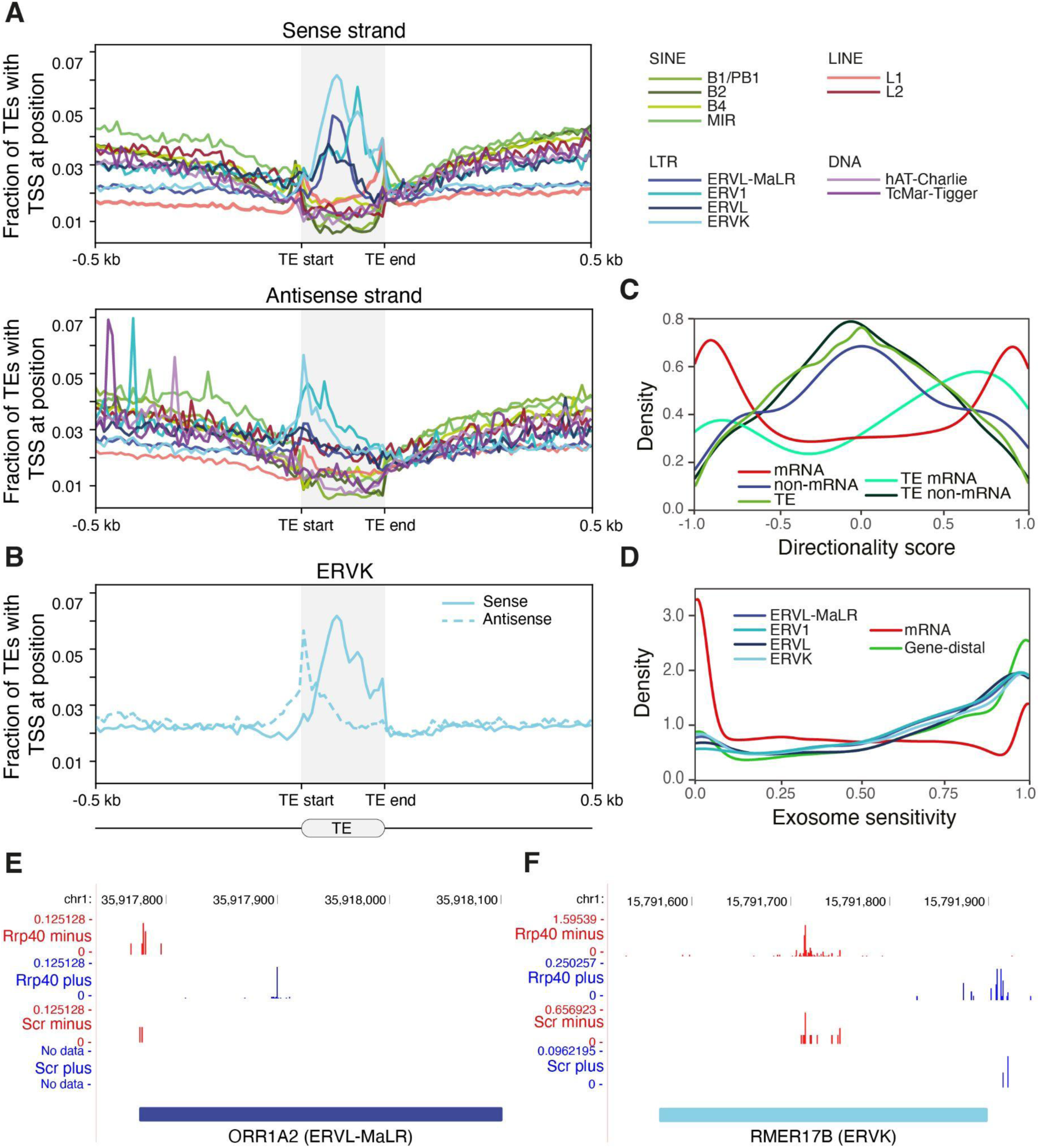
TEs are divergently transcribed into unstable RNAs. **A:** Average distribution of CAGE-inferred TSS locations (vertical axis; expression agnostic) +/- 500 bp upstream/downstream and across the body of major TE families (horizontal axis). TSS locations are visualized separately for the sense (upper panel) and antisense (middle panel) strands. **B:** Average distribution of CAGE-inferred TSS locations for the ERVK family. **C:** Transcriptional directionality score, describing the strand bias in expression levels (ranges between -1 for 100% minus strand expression and +1 for 100% plus strand expression), for mRNA and non-mRNA (non-protein-coding GENCODE transcripts) as well as TE-associated and non-TE-associated RNAs (regardless of annotation). **D:** Exosome sensitivity, measuring the relative amount of exosome degraded RNAs (ranges between 0 for RNAs unaffected by the exosome and 1 for 100% unstable RNAs), for transcripts associated with LTR families ERV1, ERVK, ERVL, and ERVL-MaLR. For comparison, exosome sensitivity is shown for mRNAs and gene-distal loci. **E-F:** Genome browser tracks for two loci of unannotated transcripts with characteristic divergent expression patterns falling on TE insertions of ORR1A2 (ERVL-MaLR; **E**) and RMER17B (ERVK; **F**) subfamilies. Pooled replicate CAGE expression levels in control (Scr) and after exosome depletion (Rrp40) split by plus (blue) and minus (red) strands are shown. For visibility reasons, the scales of CAGE signals differ between strands and conditions.

The average profiles of TSS locations for ERVs and L1 LINEs (Fig. 1A,B) indicated divergent transcription initiation, reminiscent of that of gene promoters and gene-distal enhancers (Kim et al. 2010; Andersson et al. 2014a, 2014b; Chen et al. 2016). To investigate the functional relevance of individual genomic TE insertions, we quantified transcriptional directionality in TE-associated open chromatin loci, as measured by DNase I hypersensitive sites (DHSs). The large majority of TE-associated CAGE-derived TSSs were proximal to DHSs (Supplementary Fig. 7) and the majority of TE-associated DHSs displayed balanced bidirectional transcription initiation (Fig. 1C), a hallmark of gene-distal regulatory elements with enhancer activity (Andersson et al. 2014a). This suggests that some of the investigated TEs may act as enhancers.

Divergent transcripts from enhancers and gene promoters are frequently associated with nuclear decay by the RNA exosome (Preker et al. 2008; Ntini et al. 2013; Andersson et al. 2014b). Comparing CAGE data from exosome-depleted mESCs with wildtype mESCs (scrambled shRNA control) (Lloret-Llinares et al. 2018) confirmed that TE-derived RNAs are also degraded by the exosome (Fig. 1D; Supplementary Figs. 2-4,8; Supplementary Table 1), as exemplified by CAGE data at genomic loci containing insertion sites for ORR1A2 (ERVL-MaLR) and RMER17B (ERVK) (Fig. 1E-F). The exosome sensitivity of TE-derived RNAs was similar to those of gene-distal loci, in contrast to mRNAs which are mostly protected against decay by the exosome (Fig. 1D) (Preker et al. 2008; Ntini et al. 2013; Andersson et al. 2014b). Across TE families, we generally observed more TE-associated TSSs in exosome-depleted mESCs compared to wild type mESCs (73,246 versus 28,215). Overall, more TEs with measurable transcription initiation (> 1 CAGE tag, hereafter referred to as transcribed TEs) were detected in exosome-depleted compared to wildtype mESCs (26,079 in wildtype mESCs; 64,299 in exosome-depleted mESCs; 82,383 in pooled CAGE data of exosome-depleted and wildtype mESCs). Exosome-depleted cells further displayed an increased expression level of TE-derived RNAs (Supplementary Fig. 2; Supplementary Table 1). Together, these results indicate that, although a prominent number of TEs are transcribed, the majority of derived RNAs are at least partially degraded.

Taken together, our results demonstrate that transcribed TEs in mESCs are associated with open chromatin regions accommodating divergent transcription initiation of RNAs that are substrates for endonucleolytic decay. These properties imply the potential for TEs to function as enhancers.

### Transcription of dormant ERVs reveals co-opted regulatory elements

The characteristics of transcribed TEs and the similarities between TE-derived RNAs, eRNAs and PROMPTs led us to investigate further similarities with transcribed regulatory elements. Genomic annotation of transcribed TEs in mESCs revealed that a substantial fraction was either located in gene-distal intergenic regions or overlapped with gene promoters (Fig. 2A). Across all transcribed TE families, ERV and L1 families contained the biggest fractions of TEs in intergenic regions localized at least 10 kb from the nearest gene, indicating their preference for gene-distal regulatory elements (50.6% of ERVs on average across ERV families and 59.8% of L1 LINEs). In contrast, other TE families displayed an elevated proportion of expressed TEs in genic regions. Many CAGE-inferred TSSs of transcribed TEs overlapped with open chromatin regions as defined by DHSs (62,514, ∼65%, of transcribed TEs overlapped via their TSSs with 7,431 DHSs). This suggests that a prominent fraction of transcribed TEs may act, alone (34% of DHSs overlapped TSSs of one transcribed TE) or in combination with other transcribed TEs (66% of DHSs overlapped TSSs of multiple transcribed TEs), as gene regulatory elements, but may also reflect a selfish TE tropism for open chromatin regions (Dewannieux et al. 2004; Brady et al. 2009; Jacques et al. 2013; Vierstra et al. 2014).

**Figure 2:**
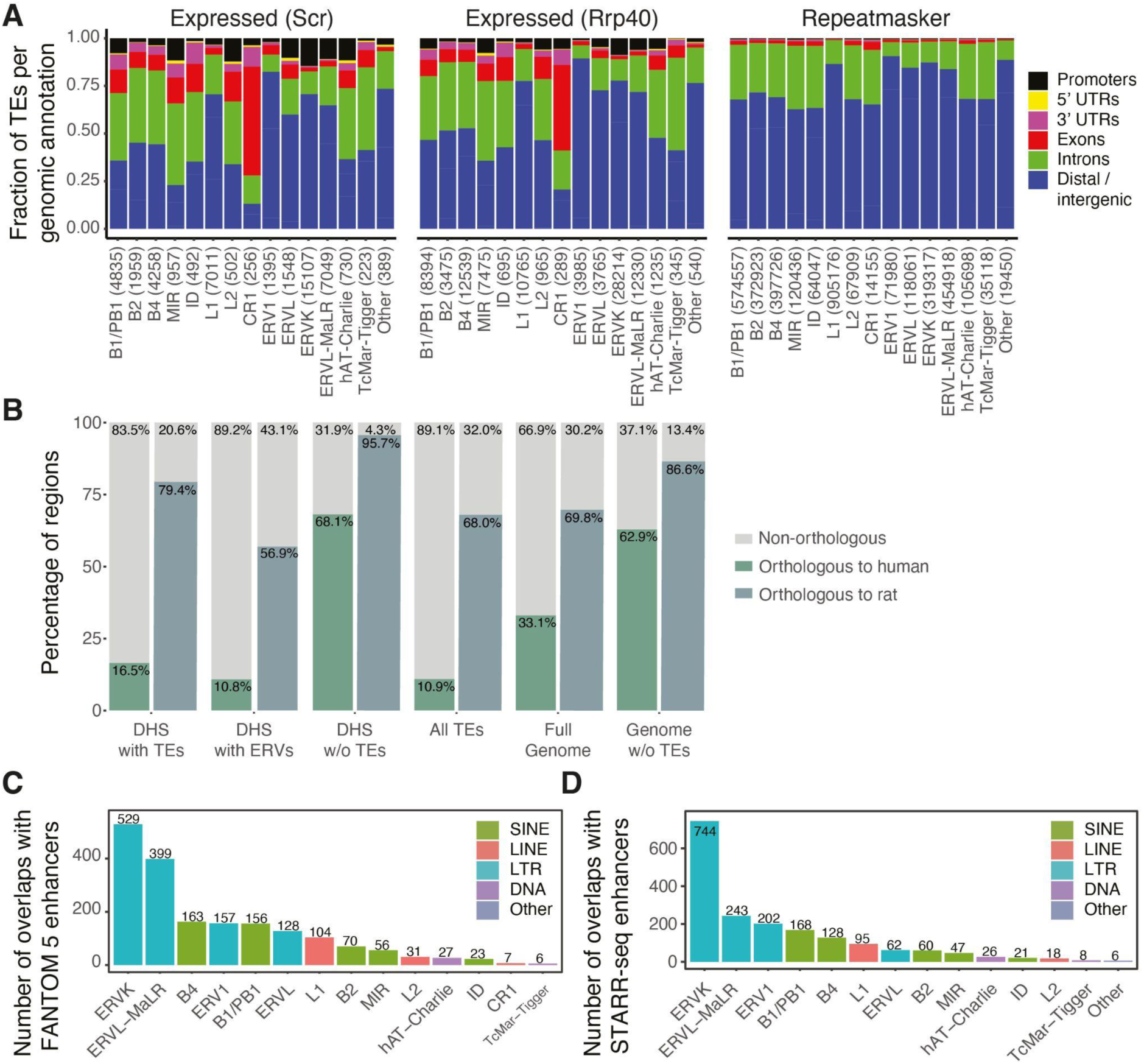
Transcription of transposable elements reveals co-opted regulatory elements. **A:** Fraction of expressed TEs in control (Scr) and exosome KD (Rrp40) mESCs as well as all TEs annotated in Repeatmasker (expression agnostic) per genomic annotation group for each TE family. The number of instances of each TE family is shown in parenthesis. **B:** Percentages of orthologous TE-associated, ERV-associated, and non-TE-associated expressed DHSs, as well as background regions between mouse and rat or human genomes. **C:** The number of transcribed TEs overlapping FANTOM 5 mouse enhancers at the TE family level. TE subfamily counts are displayed in Supplementary Figure 9A. **D:** The number of transcribed TEs overlapping STARR-seq mESC enhancers at the TE family level. TE subfamily counts are displayed in Supplementary Figure 9B.

In agreement with an association of TEs with species-specific DHSs (Vierstra et al. 2014), we observed that the majority of transcribed TE-associated DHSs in mESCs are rodent- or mouse-specific (Fig. 2B). Only 16.9% of DHS sequences had human orthologous sequences while 79.4% were orthologous to the rat genome, indicating that they derive from TEs that have accumulated after the primate-rodent split. In contrast, transcribed DHSs devoid of TEs were more conserved. Interestingly, transcribed DHSs associated with ERV subfamilies were even more mouse specific (Fig. 2B; Supplementary Fig. 10), in agreement with previous reports (Jacques et al. 2013; Sundaram et al. 2014; Miao et al. 2020), suggesting regulatory innovation through TE exaptation and that these elements contribute to the high evolutionary turnover of enhancers.

To generally assess whether transcribed dormant TEs have been co-opted as transcriptional regulatory elements, we first evaluated their association with FANTOM enhancers (Andersson et al. 2014a; Arner et al. 2015; Dalby et al. 2018), an extensively validated set of regulatory elements with predicted enhancer function inferred from bidirectional transcription initiation across a large number of cell types and tissues. Notably, 22.1% of FANTOM mouse enhancers with detectable expression in mESCs (1,979 out of 8,942) overlapped with transcribed TE-associated DHSs in mESCs (1,504 out of 7,431, 20.2%; p=1e-5, Fisher’s exact test). This indicates that TEs contribute to a sizable fraction of regulatory elements with enhancer activity. Moreover, half of this set was composed of DHSs with LTR elements, in particular ERVKs and ERVL-MaLRs (Fig. 2C; Supplementary Fig. 9A), consistent with previously reported bidirectional transcription of ERVs (Fort et al. 2014). Of note, given that FANTOM enhancers were neither identified from exosome depleted mESCs nor multi-mapping rescued reads, it is conceivable that some TE-associated enhancers are not present in the FANTOM enhancer set.

Since FANTOM enhancers were predicted based on the same property that we had observed for transcribed TEs, namely divergent transcription initiation, alternative experimental data is required to evaluate the enhancer potential of TEs (Halfon 2019). To this end, we considered the *in vitro* enhancer potential of transcribed TEs using genome-wide mESC STARR-seq data (Peng et al. 2020). 18.9% of the transcribed TE-associated DHSs (1,404 out of 7,431) overlapped with open chromatin-associated STARR-seq enhancers (1,949 out of 7,078, 26.2%; Fig. 2D; Supplementary Fig. 9B). Again, LTRs, in particular ERVKs, constitute a sizable fraction of these enhancers, in agreement with the FANTOM enhancer overlap. In contrast, transcribed L1 LINE elements, despite frequently residing in gene-distal intergenic loci (Fig. 2A), rarely overlapped with FANTOM or STARR-seq enhancers. L1 elements are therefore less likely to act as enhancers in ESCs, in line with human LINEs (Barakat et al. 2018). Interestingly, in agreement with the strong association between transcription initiation and *in vitro* enhancer potential (Andersson et al. 2014a; Wu et al. 2014; Rennie et al. 2018), we observed that non-transcribed TE-associated DHSs were to a lesser degree validated by STARR-seq enhancers than transcribed ones (p<2e-16 for ERVKs and ERVL-MaLR, Fisher’s exact test).

Combined, the STARR-seq and FANTOM sets indicate that 2,378 (∼32%) transcribed TE-associated DHSs in mESCs are likely enhancers. Example loci are displayed in Figure 3, which clearly illustrate the association between divergent transcription initiation and enhancer activity from individual TEs (Fig. 3A; RLTR41 insertion) or, in some cases, pairs of TEs (Fig. 3B; RLTR41 in pairs with MYSERV-int and RMER10B insertions).

**Figure 3:**
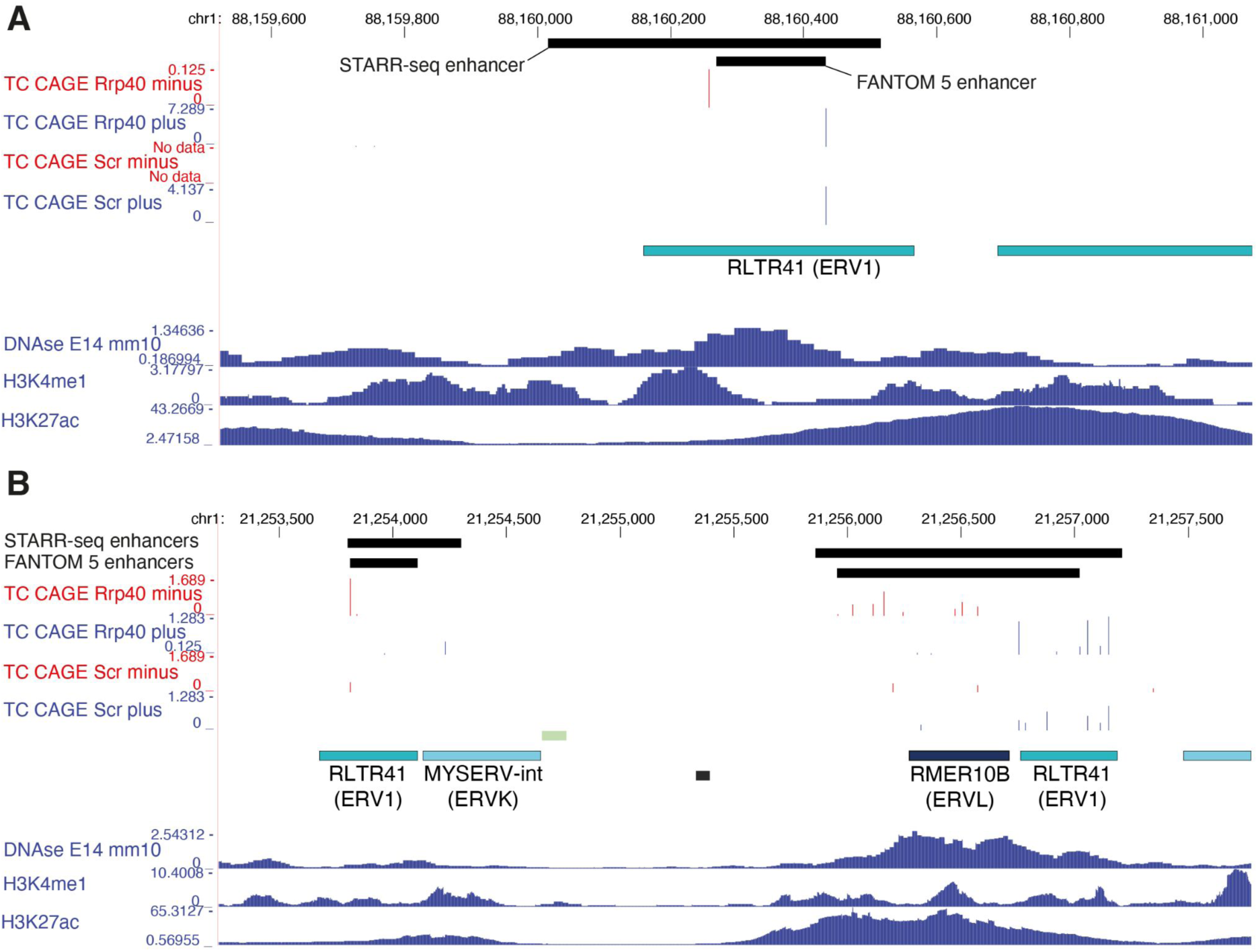
TE insertions co-opted as divergently transcribed enhancers. **A-B:** Genome browser tracks for two intergenic loci showing TPM-normalized CAGE data pooled across replicates and split by plus (blue) and minus (red) strands. Shown are also the locations of FANTOM5 mouse enhancers and STARR-seq mESC enhancers and signal tracks for ENCODE DNase-seq data and H3K4me1 and H3K27ac ChIP-seq data for E14 mESCs. The CAGE signals identify divergent transcription initiation from ERV1 RLTR41 insertions, alone (**A**) and in pairs (**B**) with TE insertions of RMER10B and MYSERV-int.

### Transcribed TEs carry chromatin features of regulatory elements

The strong overlap of ERVK elements and weak overlap of L1 elements with FANTOM and STARR-seq enhancer sets implies differences in the regulatory potential among TE families (Fig. 2C,D). To investigate these differences further, we assessed the chromatin states at transcribed TEs. For this, ChIP-seq signal for histone modifications was aggregated around CAGE-inferred TSSs in TE-associated DHSs (The ENCODE Project Consortium 2012; Yue et al. 2014). Hierarchical clustering of these aggregated signals revealed six major chromatin states (Fig. 4A; Supplementary Fig. 11).

**Figure 4:**
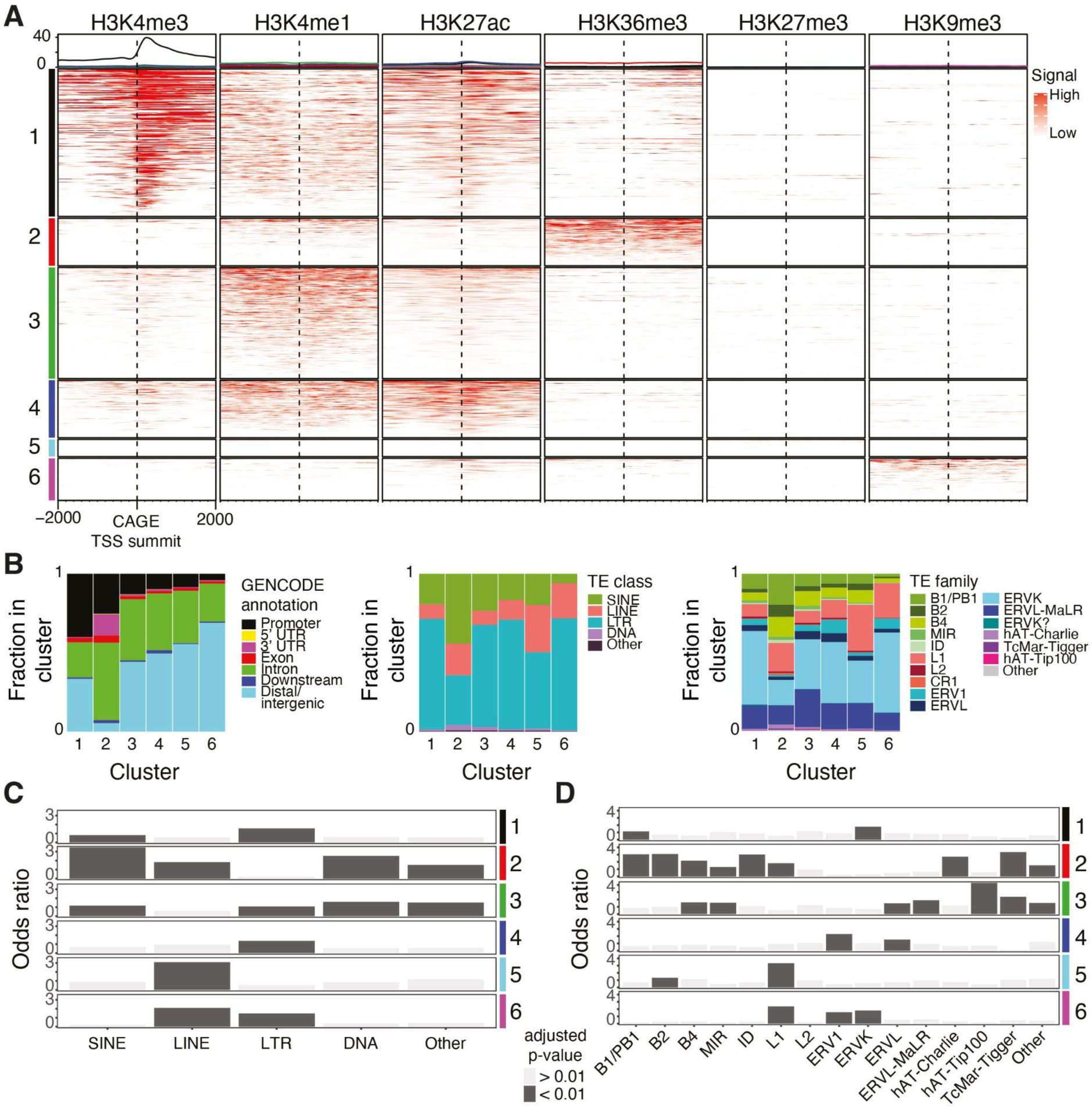
Transcribed TEs exhibit chromatin features of regulatory elements. **A:** Hierarchical clustering of histone modification (ChIP-seq) signals +/- 2,000 bp around the summits of TE-associated clusters of CAGE-inferred TSSs (CAGE tag clusters, Methods). The ChIP-seq signal is shown as fold-change over input control. Clusters are represented in rows (color coded in left legend) and histone modifications in columns. Average distributions of ChIP-seq signals for each cluster are shown (top panel). **B:** Annotations of TE-associated CAGE tag clusters based on GENCODE and RepeatMasker TE classes and families. **C,D:** Bar plots of odds ratios of enrichments (Fisher’s exact test) of TE classes (**C**) and TE families (**D**) in each cluster.

The majority of loci (clusters 1, 3, 4) carried histone modification signal associated with active regulatory elements (Creyghton et al. 2010; Heintzman et al. 2007; Robertson et al. 2008): H3K4me1/3 and/or H3K27ac (Fig. 4A; Supplementary Fig. 12). The elements in cluster 3 and 4 were mainly intronic or gene-distal (>80%; Fig. 4B) and displayed H3K4me1 signal, associated with weak transcription (Core et al. 2014; Andersson and Sandelin 2020), in combination with H3K27ac. We observed a strong association with LTRs (Fig. 4B,C), in particular ERVs (Fig. 4B,D), in these two clusters, reinforcing the association statistics with STARR-seq and FANTOM enhancers. Although cluster 3 loci were associated with many TE families, a considerable fraction of TEs belong to the ERVL, ERVL-MaLR and B4 TE families (ERVL: 871, 33.9% of clustered ERVLs; ERVL-MaLR: 4,106, 37.4% of clustered ERVL-MaLRs; B4: 1,597, 35.5% of clustered B4s). ERV1s and ERVLs were enriched among cluster 4 TEs (Fig. 4D). In addition, transcribed STARR-seq associated DHSs displayed comparable expression levels and similar histone modifications at TE and non-TE loci (Supplementary Fig. 13), suggesting that TE-derived enhancers operate at similar activity levels as non-TE enhancers.

A considerable fraction of TE insertions (cluster 1) displayed strong H3K4me3 signal, associated with promoter activity of highly expressed transcription units, e.g., mRNA genes (Core et al. 2014; Andersson and Sandelin 2020). The majority of these fell close to GENCODE annotated TSSs and were enriched with SINE (B1 and proto-B1) and LTR (ERVK) elements (Fig. 4B-D). Hence, TEs are not only evolutionary co-opted into distal regulatory elements, but also gene promoters. Indeed, out of 7,431 transcribed TE-associated DHSs, 1,012 (13,6%) were associated with GENCODE annotated mRNA gene TSSs and 558 (7,5%) with GENCODE long non-coding RNA (lncRNA) gene TSSs. Complementary chromatin state segmentation analysis confirmed the strong bias of transcribed TEs having histone modifications indicative of active regulatory elements (Supplementary Fig. 14).

Interestingly, we found some ERV1s and ERVKs, in addition to L1 LINEs, to be associated with both H3K27ac and H3K9me3 (cluster 6). The latter is present at repressed TEs in heterochromatin (Rowe et al. 2013; Matsui et al. 2010; Maksakova et al. 2013; Karimi et al. 2011) and the combination of both repressing (H3K9me3) and activating (H3K27ac) histone modifications has been suggested to keep these TE-associated regulatory elements in a partially activated state (He et al. 2019). We also identified a group of transcribed TE loci (cluster 5) that was highly enriched with L1s but overall, only carried low levels of histone modifications. Some of these loci were associated with H3K27me3 signal, indicative of polycomb-mediated repression. TE loci of cluster 5 and 6 thus likely represent repressed regulatory elements with low transcriptional activity, which may facilitate later full activation, although we cannot rule out that this is an effect of mESC clonal heterogeneity.

Taken together, through several lines of evidence we demonstrate that dormant TEs, in particular endogenous retroviral elements, have frequently been repurposed into regulatory elements with enhancer and promoter activities in mESCs.

### The TF binding repertoires and regulatory topologies of ERV subfamilies indicate involvement in distinct regulatory programs

TF binding to regulatory elements is the key determinant of regulatory activity and the basis of cell-type specificity. To assess how TEs co-opted as regulatory elements may contribute to transcriptional regulatory programs in mESCs, we performed a TF binding site enrichment analysis of transcribed DHSs associated with TE insertions from 224 LTR subfamilies versus all genomic loci of CAGE-inferred TSSs in mESCs (Methods). This analysis thus reveals TFs that are enriched or depleted in TEs compared to what we would expect from regulatory elements in general. Interestingly, predicted binding sites for several TFs, including pluripotency factors Nanog, Sox2 and Oct4, were enriched in transcribed LTRs compared to non-TE associated TSSs (Benjamini-Hochberg FDR-adjusted p<1e-3). Binding sites for these TFs were further found in a large number of LTR insertions (in 70.9%, 24.5% and 11,5% of all 19,436 transcribed DHS-associated LTRs versus in 40.3%, 9,6% and 5.2% of CAGE-inferred TSSs for Nanog, Sox2 and Oct4, respectively). These results indicate that LTRs are a major source of regulatory elements controlled by pluripotency factors in mESCs. These results are in line with those observed for human ESCs (Kunarso et al. 2010; Barakat et al. 2018), despite the low conservation of LTRs between species (Fig. 2B) (Glinsky and Barakat 2019).

Given their strong association with regulatory elements, we focused on putative TF binding sites in DHSs of transcribed TEs across ERV subfamilies. In fact, we observe a variability in enrichment of various TF binding site sequences in specific ERV subfamilies indicating diverse regulatory potentials of TEs and specific TF-TE- associations (Fig. 5A; Supplementary Fig. 15A; Supplementary Table 2). We observed a positive correlation between enrichments of binding sites for pluripotency factors, e.g., Sox2 and Esrrb (Pearson’s r = 0.54), as well as Sox2 and Nanog (Pearson’s r = 0.47; Fig. 5B; Supplementary Fig. 15B). Differences in enrichments for Sox2 and Nanog were, however, found for some ERV subfamilies, as seen for instance for ORR1A1 and ORR1A2 (Fig. 5A), indicating that calculated motif enrichment similarities between these two TFs are not necessarily driven by motif similarities. The overall enrichments of Sox2, Nanog and Esrrb generally agreed with Isl1, Bcl6 and Oct4 (Fig 5B; Supplementary Fig. 12B), indicating that certain ERV subfamilies carry binding sites for a wide variety of pluripotency factors. Binding site sequences for all these TFs and the core pluripotency factor Oct4 were enriched in RLTR41, RLTR23, MMERVK9C_I_int, MLTR25A, and RLTR11A (mean log_2_ enrichment > 1).

**Figure 5:**
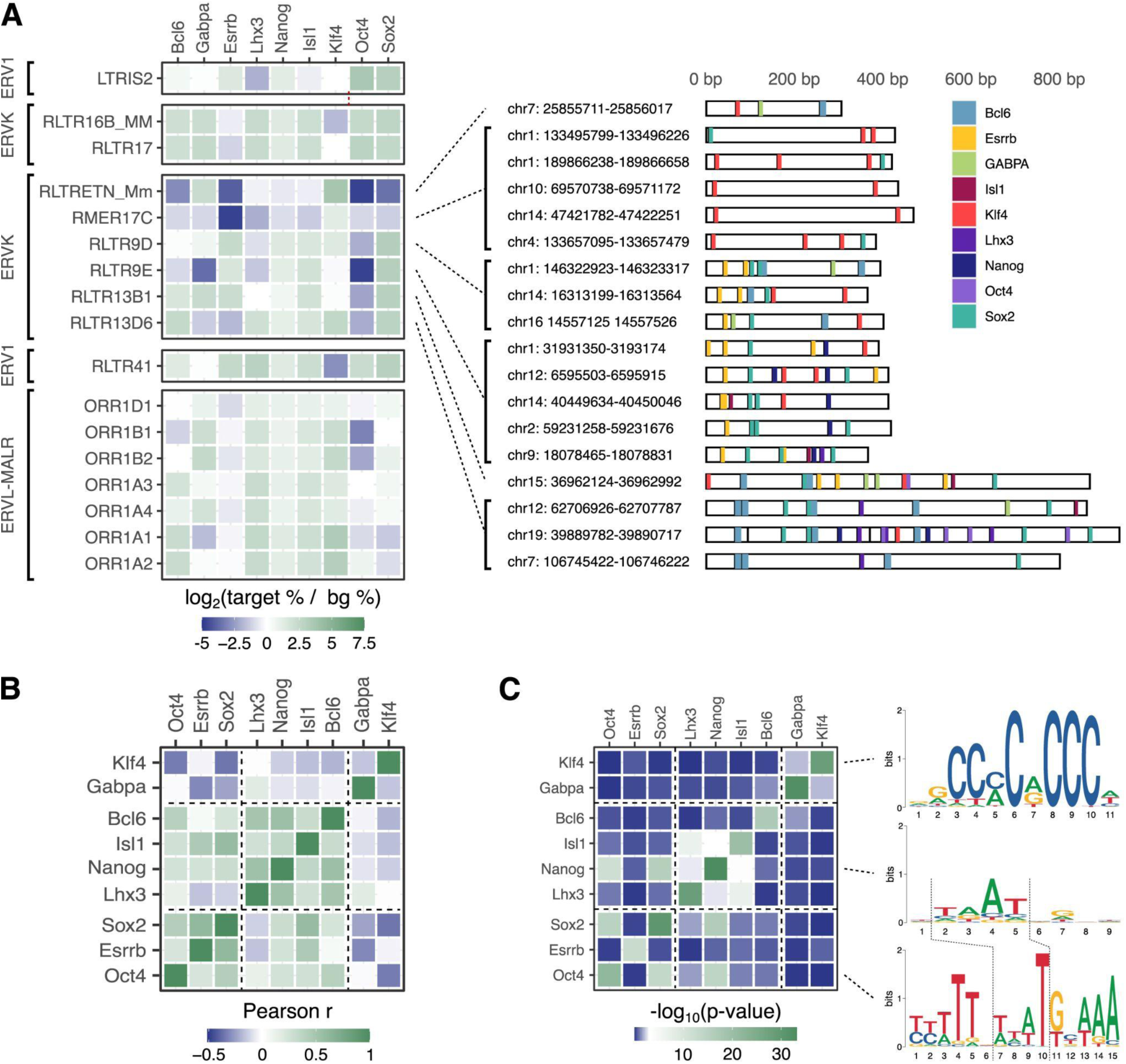
ERV subfamilies contribute to distinct enrichments of binding sites for pluripotency factors. **A:** Motif enrichments for selected TFs (columns) in transcribed TEs across selected ERV subfamilies (rows) versus a background of non-TE genomic regions +/- 200 bp around the summits of all CAGE-inferred TSS clusters. White cells indicate either no enrichment or non-reported values. The full TF enrichment heatmap is shown in Supplementary Figure 15A. Example genomic insertion sites for ERVK subfamilies are shown to illustrate their differences in carrying putative TF binding sites for Sox2, Esrrb and Oct4. **B:** Correlations (Pearson’s r) between TF motif enrichments across all ERV subfamilies (as shown in Supplementary Fig. 15A). The full heatmap of correlations is given in Supplementary Figure 15B. **C:** Similarity (q-value) between binding motifs for selected TFs. Sequence logos (right) for Oct4, Nanog and Klf4 exemplify differences and similarities of TF binding sites.

We noted that SOX family related motifs, including that for Sox2, were enriched in specific ERV subfamilies (most notably in RLTR9D, RLTR9E, LTRIS2, RLTR13B1, RLTR41). Similarly, we identified a specific enrichment of KLF factors, including Klf4 (most highly enriched in RLTRETN_Mm, ORR1A1, ORR1A2, and RLTR10A). ERVL-MaLRs ORR1A1 and ORR1A2 were, in addition to Klf4, also enriched with Nanog, Isl1, Bcl6, and Lhx3, but interestingly not Esrrb and Oct4 binding sites. This suggests that, by carrying different repertoires of TF binding sites, ERV subfamilies may contribute differently and distinctly to the pluripotency regulatory program (Supplementary Fig. 15). Similarly, Lhx3 binding sites co-occurred with those of Esrrb in RLTR41 ERV1s, but were depleted from Esrrb-enriched ERVKs RLTR9D and RLTR13B1. In addition, the top enriched ERVs for Oct4 binding sites (LTRIS2, RLTR17, RLTR16B_MM, RLTR41) were depleted from those of Klf4.

The specific enrichments or depletions of Klf4 or Oct4 binding sites in certain ERV subfamilies indicate that the Klf4 and Oct4 ERV-derived regulatory networks have partially evolved independently from those of Sox2 and Nanog. This is consistent with distinct binding preferences (Fig. 5C) and context-dependent cooperativity between Sox2 and Oct4 (Li et al. 2019), which is limited during pluripotency maintenance (Malik et al. 2019). We note that the binding preferences for Oct4 and Nanog could allow for the derivation of new binding sites for one TF from ancestral binding site sequences of the other through mutations (Fig. 5C). However, the dissimilarity of the Klf4 motif with those of Oct4 and Nanog indicates that evolutionary acquisition of Klf4 binding sites likely requires new transposition events.

In addition to the enrichment of pluripotency TFs, we identified putative ERV-derived binding sites for a broad range of TFs (Supplementary Fig. 15A). For instance, binding site sequences for ETS factors, involved in a large variety of gene regulatory programs and across cell types, were highly enriched among ERVL-MaLR subfamilies ORR1E and ORR1D2, which were further depleted of TF binding sites for Oct4 and Sox2. This suggests that transposition of TF binding sites native to LTR retrotransposons or the subsequent birth of new TF binding sites through mutations (Sundaram et al. 2017) could impact several regulatory programs, including those distinct from the naive pluripotency.

To further investigate the regulatory programs contributed to by ERVs co-opted as regulatory elements, we linked each gene-distal, transcribed ERV-associated DHS with its predicted target gene from activity-by-contact (ABC) modeling (Fulco et al. 2019). The resulting gene ontology enrichments of linked genes indicate a highly specific association of individual ERV subfamilies with gene regulatory programs (Fig. 6A; Supplementary Fig. 16), including regulation of genes involved in immunity by RLTR13B1 elements and regulation of genes involved in transcription, specifically basal TFs in the TFIID complex, by MLTR14 elements (Fig. 6B).

**Figure 6:**
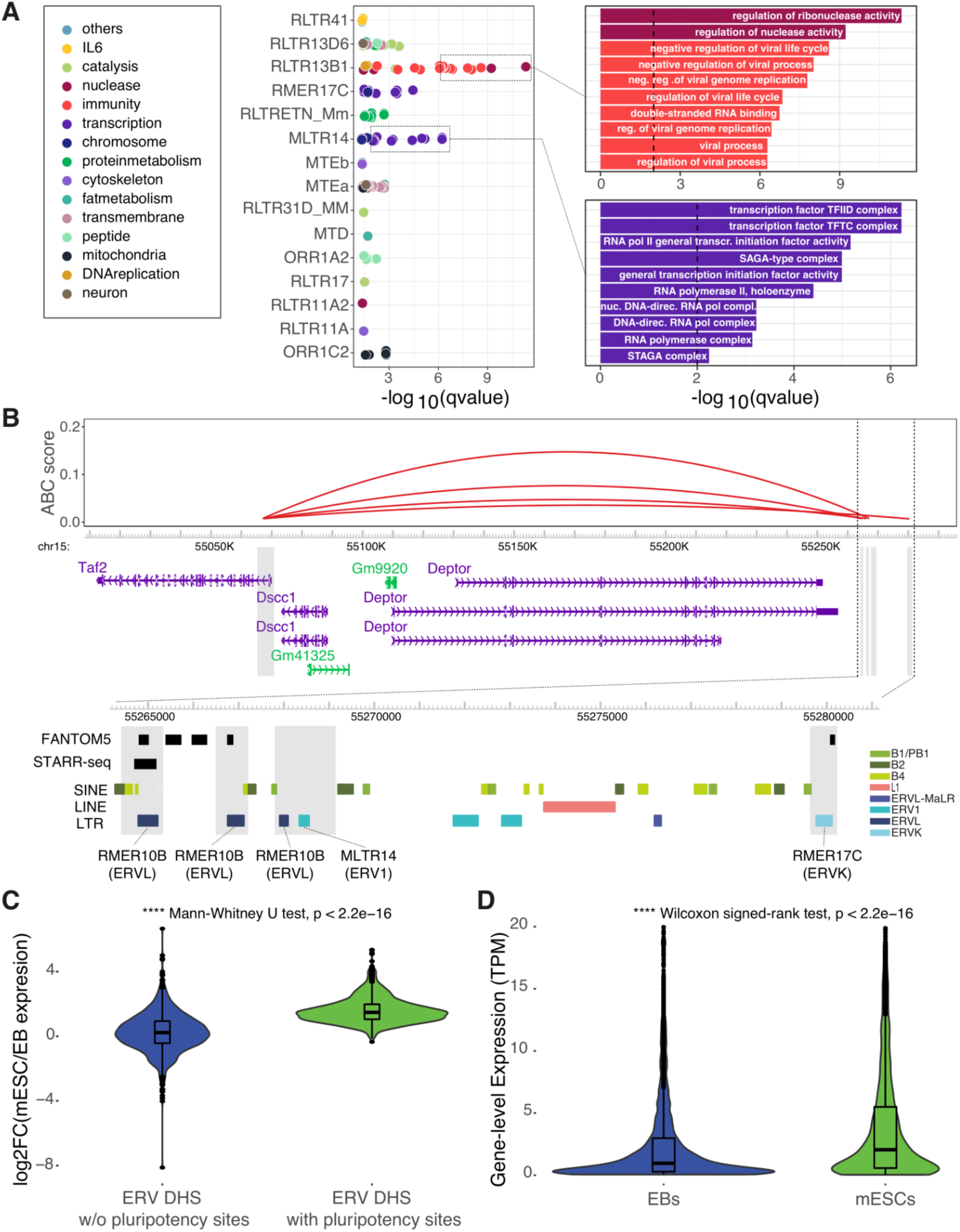
ERV subfamilies contribute to distinct gene regulatory programs. **A:** GO term enrichment for putative target genes (ABC) of gene distal ERVs split by ERV subfamily (foreground) versus all ABC-predicted target genes (background). For ease of visualization, gene ontology terms are colored by manually curated process or function, and the underlying gene ontology term enrichments are shown for RLTR13B1 and MLTR14. Full results are provided in Supplementary Figure 16. **B:** Predicted enhancer interactions with the promoter of gene Taf2 (TATA-Box Binding Protein Associated Factor 2). The four enhancers are marked by grey boxes and a zoom-in is provided below, showing overlaps with ERV insertions of RMER10B, MLTR14 and RMER17C. Tracks for GENCODE (M19) transcripts, FANTOM5 enhancers, STARR-seq enhancers, and TEs provided by RepeatMasker are shown. **C:** Expression fold-change (log_2_), as measured by CAGE, in mESCs versus EBs at ERV-associated DHSs that carry (right) or not (left) predicted binding sites for pluripotency factors. **D:** Gene-level expression (TPM normalized), as measured by CAGE, quantified for ABC-linked genes of transcribed TE-associated DHSS that carry predicted binding sites for pluripotency factors.

The enrichment of pluripotency factors in transcribed ERV-associated DHSs suggests that the regulatory activities of these elements have a specificity towards cell types in which these TFs are active. To investigate their putative cell-type restricted activity, we quantified their expression using CAGE data for exosome-depleted samples derived after 3 days differentiation of mESCs into embryoid bodies (EBs) (Lloret-Llinares et al. 2018). In agreement with their predicted role in mESCs, the expression of ERV-associated regulatory elements with inferred binding sites for pluripotency TFs (as given in Fig. 5A) were generally reduced in EBs, while those not containing such sites displayed a similar expression level in mESCs and EBs (Fig. 6C). Accordingly, genes whose ABC-linked ERV-associated enhancers contained binding sites for pluripotency factors displayed lower expression in EBs (Fig. 6D). These observations reflect a reduced activity of pluripotency TFs in EBs and demonstrate that expression of ERV-associated DHSs can be used as a marker for their cell-type specificity in regulatory activity.

Taken together, our results indicate that ERV-derived regulatory elements transcribed in mESCs contribute in a specific manner to the pluripotency regulatory network through their binding sites for pluripotency TFs. Although we identified putative TF binding sites for multiple TFs for each ERV subfamily, the TF enrichments across ERV subfamilies were highly distinct (Fig. 5; Supplementary Fig. 12). Together with the diversity of TF binding sites identified (Supplementary Fig. 12A) and the diversity of functions of target genes (Supplementary Fig. 13), this suggests that each ERV subfamily contributes to distinct regulatory programs and pathways.

## Discussion

Transcriptional regulation of ESCs is multi-faceted. On one hand, regulatory elements act as binding platforms for transcription factors controlling the transcriptional activities of genes involved in activities not necessarily specific to ESCs, such as metabolism, transcription, stress response and replication. In parallel, ESCs must maintain plasticity to ensure potent differentiation capabilities. At the core of such activities are highly specialized and conserved TFs, including Oct4, Sox2 and Nanog, which, through targeted transcriptional regulation, maintain a naive pluripotent state. While gene regulation in ESCs by these TFs is highly conserved across Metazoa, their respective binding sites are not. TEs have been suggested to contribute to stabilizing the gene regulatory functions of TFs by providing regulatory sequences with the required binding sites.

We here demonstrate that retrotransposons contribute to a sizable fraction of such regulatory innovation. Using an accurate and unbiased approach based on genome-wide profiling of TSSs, we systematically investigate the regulatory potential and transcriptional activities of the wide repertoire of mouse TEs in mESCs. We show that a fraction of TEs are transcribed and that these display balanced divergent transcription initiation patterns within sites of open chromatin. We provide, to the best of our knowledge, the first evidence that retrotransposon-derived RNAs are targeted by the nuclear exosome for degradation. As many as a third of divergently transcribed TE-associated DHSs are supported by *in vitro* enhancer potential derived from STARR-seq data or by overlap with, rigorously validated, previously predicted enhancer sets from FANTOM. In addition, we find a large overlap with annotated gene promoters, demonstrating that at least a half of transcribed TE-associated DHSs in mESCs are regulatory elements (enhancers or promoters). Transcription start site profiling by CAGE thus lends itself as an accurate approach for genome-wide surveys of the regulatory activity of TEs, for which predictions based on open chromatin and histone modifications yield considerably lower validation rates (Todd et al. 2019; Barakat et al. 2018). We observed that a substantial fraction of transcribed TE-derived regulatory elements were non-orthologous to rat or human genomes, demonstrating a high degree of regulatory innovation by TEs. Our TSS-centric approach thus allows for an unbiased, systematic investigation of the regulatory potential across TE subfamilies, beyond a selected few subfamilies.

We note that disagreements in validation rates to a recent study (Todd et al. 2019) could further be due to difference in validation assays used. While CRISPR interference (CRISPRi), used by Todd et al. (Todd et al. 2019), has the potential to better reflect *in vivo* activities, the false-negative rate of CRISPRi remains a concern (Gasperini et al. 2020). In addition, redundant enhancers may buffer regulatory effects (Hong et al. 2008; Cannavò et al. 2016). Therefore, an enhancer could still have a causal role in gene regulation even though no observable effect can be measured by CRISPRi. As such, *in vitro* measurements, e.g., derived from STARR-seq, are therefore better suited to reveal the enhancer potential of a regulatory element, even though such an approach may suggest enhancers that are not active *in vivo*. Further studies are necessary to properly compare the quantified activities of regulatory elements between STARR-seq and CRISPRi.

Endogenous retroviral elements were most frequently transcribed, and ERVKs stand out as the largest contributor of regulatory elements with enhancer potential in mESCs. This bias is likely explained by their general enrichment of binding sites (Bourque et al. 2008; Sundaram et al. 2017) and binding site sequences (Fig. 5A) for TFs regulating naive pluripotency, including Oct4, Sox2 and Nanog. However, we do acknowledge that mapping of sequencing reads might skew our focus to evolutionary older ERV subfamilies (Simonti et al. 2017), which have accumulated more mutations than younger ones and therefore have a higher chance of mapping reads uniquely (Jin et al. 2015). However, any mapping bias cannot explain differential TF enrichments across ERV subfamilies or their cell-type specific expression patterns.

The diverse TF enrichments observed across ERV subfamilies indicate a major contribution of ERVs to the landscape of regulatory elements and a wide variety of gene regulatory programs. Interestingly, we observed varying degrees of over-representation of putative TF binding sites in ERV subfamilies, including those enriched with the wide repertoire of pluripotency TFs, those partially depleted of specific pluripotency TFs, and those enriched with non cell-type specific TFs, like the ETS superfamily. We noted a general strong co-occurrence of binding site sequences for Nanog, Sox2, and Esrrb, while Oct4 and Klf4 were depleted in some subfamilies. While some of these TFs have similar motifs, e.g., Nanog, Sox2, and may therefore overestimate co-occurrences, the specific enrichments and depletions argue for unique contributions by ERV subfamilies to the regulatory programs of mESCs, which is confirmed by the specific functions of putative target genes by ERVs predicted to be co-opted as gene-distal enhancers. Further, our data suggest that the regulatory landscape of naive pluripotency in mESCs has been shaped independently by multiple subfamilies of ERVs.

In agreement with previous work (Bourque et al. 2008; Kunarso et al. 2010; Barakat et al. 2018), we observed a substantial enrichment of binding sites for pluripotency factors in ERV-associated regulatory elements. However, our transcription-centric approach identified many ERV subfamilies with enrichment for Nanog or Oct4 not revealed when focusing on histone modifications (e.g., H3K27ac) and chromatin accessibility (Kunarso et al. 2010), including ORR1A1, ORR1A2, RMER17B, LTRIS2, RLTR9E, RMER19B for Nanog and LTRIS2, RLTR16B_MM, MLTR25A, RLTR23, BGLII_B, RMER1B for Oct4. It is thus likely that utilizing transcription initiation profiling to infer regulatory activity more specifically identifies regulatory elements for which relevant DNA sequences, e.g., binding sites for pluripotency factors, are linked with cell-type specific transcriptional and regulatory activity. This is confirmed by seemingly cell-type restricted expression of pluripotency-associated ERVs and that of their putative target genes, when expression is compared between mESCs and EBs (Fig. 6C,D).

In addition to the core pluripotency TFs, we observed an enrichment of binding sites for multiple TFs in several ERV subfamilies. Since ERVs originate from ancestral genomic insertions of retroviral elements, the ancestral ERV sequences must therefore have either carried the full repertoire of TF binding sites that have been maintained through evolution or have served as substrates for the evolutionary acquisition of diverse TF binding sites through mutations (Sundaram et al. 2017; Sun et al. 2018). That way, ERVs may offer enough sequence diversity to allow encoding for enhancer activity even in the absence of binding sites for specific master regulators of mESCs (Singh et al. 2021).

Taken together, we present a systematic characterization of the transcriptional activities, RNA decay patterns, chromatin signatures, and regulatory potential of TEs in mESCs. Our results demonstrate a sizable contribution of TEs to regulatory innovation and the regulatory landscape of naive pluripotency. We further show that expression of TE-associated open chromatin regions is indicative of cell-type restricted regulatory activity. Charting their dynamic activities over development will thus be an important next step to further our understanding of the cell-type specific roles of TEs in transcriptional regulation.

## Author contributions

R.A. and S.B conceived the project; S.B performed most analyses, with support from R.K., N.A. and M.S.; S.B., R.K. and R.A interpreted results and wrote the manuscript; all authors reviewed the final manuscript.

## Supplementary Note: evaluation of multi-mapping rescue of CAGE reads

Of all mouse TE insertion events (3,677,522), only 1.3% (46,424) were associated with uniquely mapped CAGE tags in mESCs. Expressed TEs were associated with ∼4% of all identified TSSs in mESCs and ∼2% of uniquely mapped CAGE reads. Employing the MuMRescueLite multi-mapping rescue approach (Faulkner et al. 2008, 2009; Hashimoto et al. 2009) led to an increase in the number of detected expressed TEs (the number of detected TEs increased by 77.5% to 82,383 TEs) and an overall higher expression level (Supplementary Figs. 2B, 3, 4).

Comparison of TE expression quantification after MuMRescueLite multi-mapping with an alternative quantification strategy (TELocal) (Jin et al. 2015), based on maximum likelihood alignments to annotated repeats, revealed distinct TE instances identified as expressed by each approach (Supplementary Fig. 5). The results show that MuMRescueLite and TELocal yield a comparable number of unique zero expression TE instances per family and that TE expression values for TE insertions identified by both approaches are comparable (Spearman’s rho ranging between 0.53 and 0.7; Supplementary Fig. 5), in particular for ERVs. Note that TPM values are not directly comparable since TELocal is based on a predefined set of regions, while with MuMRescueLite we considered mapping to the whole genome. However, the benefit of being able to study TSSs at base pair resolution, which is not possible with TELocal that yields one expression value per locus, and to quantify the expression levels of TSSs genome-wide not having to rely on custom annotations of TE insertions made us opt for the probabilistic multi-mapping rescue (MuMRescueLite) strategy.

We further analyzed TE-associated transcription initiation events using CAGE data from exosome-depleted HeLa cells (Andersson et al. 2014b). We observed a similar increase in TE expression levels upon exosome depletion as observed in mESCs, and that probabilistically rescued multi-mapping reads increased the number of detected transcribed TEs (Supplementary Fig. 1).

We next investigated how rescuing multi-mapping CAGE reads affects expression level quantification of genes with TE-associated promoters in HeLa cells. Using generalized linear Poisson regression, we observed that the multi-mapping rescue approach improved the agreement between promoter-derived gene expression levels inferred from HeLa CAGE (Andersson et al. 2014b) with those from HeLa RNA-seq data (Andersen et al. 2013) quantified from exonic reads (p=0.0112 and p<2e-16 for uniquely mapped CAGE data versus data also including rescued reads, respectively, F-test). In particular, we utilized RNA-seq expression summarized to the gene-level as the response variable of the generalized linear model and gene-level expression as measured by CAGE-seq using unique or rescued alignment approaches as continuous predictor variables, quantified as explained in the Methods. To better explore the ambiguity introduced in our CAGE dataset with the use of multi-mapping rescuing, several predictor variables of gene-level rescued CAGE expression with different window parameters (6, 20 and 50 base pairs around each multi-mapping read) together with the CAGE expression derived from uniquely mapped reads were tested and a backward elimination approach was used to remove predictor variables with non-statistically significant p-values. We were able to identify a significant relationship both for the unique CAGE and rescued CAGE gene-level expression with a 50bp window to the RNA-seq expression, with a higher explanatory power after using the backward elimination approach (adj. R squared=0.311 for the 2-variable model versus 0.278 for the 4-variable model; lower BIC criterion for model selection=1.219621 for the 2-variable model versus 1.631852 for the 4-variable model). This shows that quantification of TE-derived transcripts with CAGE may underestimate their abundances and that rescuing multi-mapping reads with a proper window parameter around multi-mapping reads can alleviate some of these challenges.

## Supplementary Figures

**Supplementary Figure 1.**
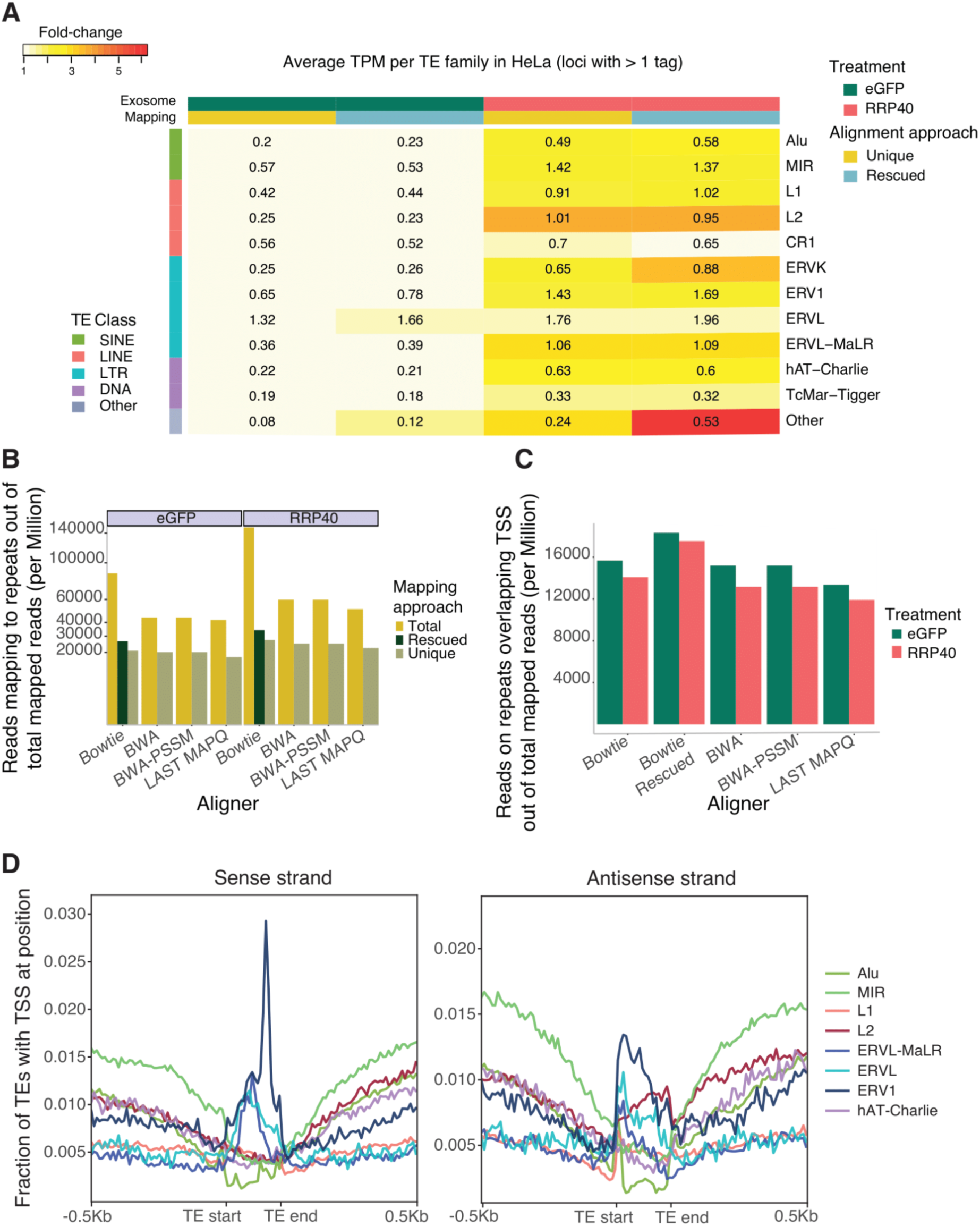
**A:** Heatmap of average TPM-normalized expression values quantified at the CTSS level for major TE families in HeLa S2 cells. Values for control (eGFP) or exosome-depleted (RRP40) CAGE libraries and using only uniquely mappable reads (Unique) or by employing the MumRescueLite algorithm (Rescued) are shown per column. The values in the cells are calculated as TPM values divided by the union of TEs expressed in at least one of the columns/conditions, thus representing a comparable average TPM value. The color key represents the fold-change versus the value for uniquely mappable reads in control samples. **B:** The number of CAGE tags mapping to TEs out of the total number of mapped reads (per million reads) using different aligners (horizontal axis) and different downstream approaches (keeping all multi-mapping reads, all uniquely mappable reads and Multimapping rescued reads with Bowtie). **C:** The number of CAGE tags mapping to TEs overlapping GENCODE TSSs out of the total number of mapped reads (per million reads) using different aligners and downstream approaches (horizontal axis) as quantified in control (eGFP) and exosome-depleted (RRP40) CAGE libraries. **D**: Average distribution of CAGE-inferred TSS locations in HeLa S2 cells (vertical axis; expression agnostic) +/- 500 bp upstream/downstream and across the body of major TE families (horizontal axis). TSS locations are visualized separately for the sense (left panel) and antisense (right panel) strands.

**Supplementary Figure 2.**
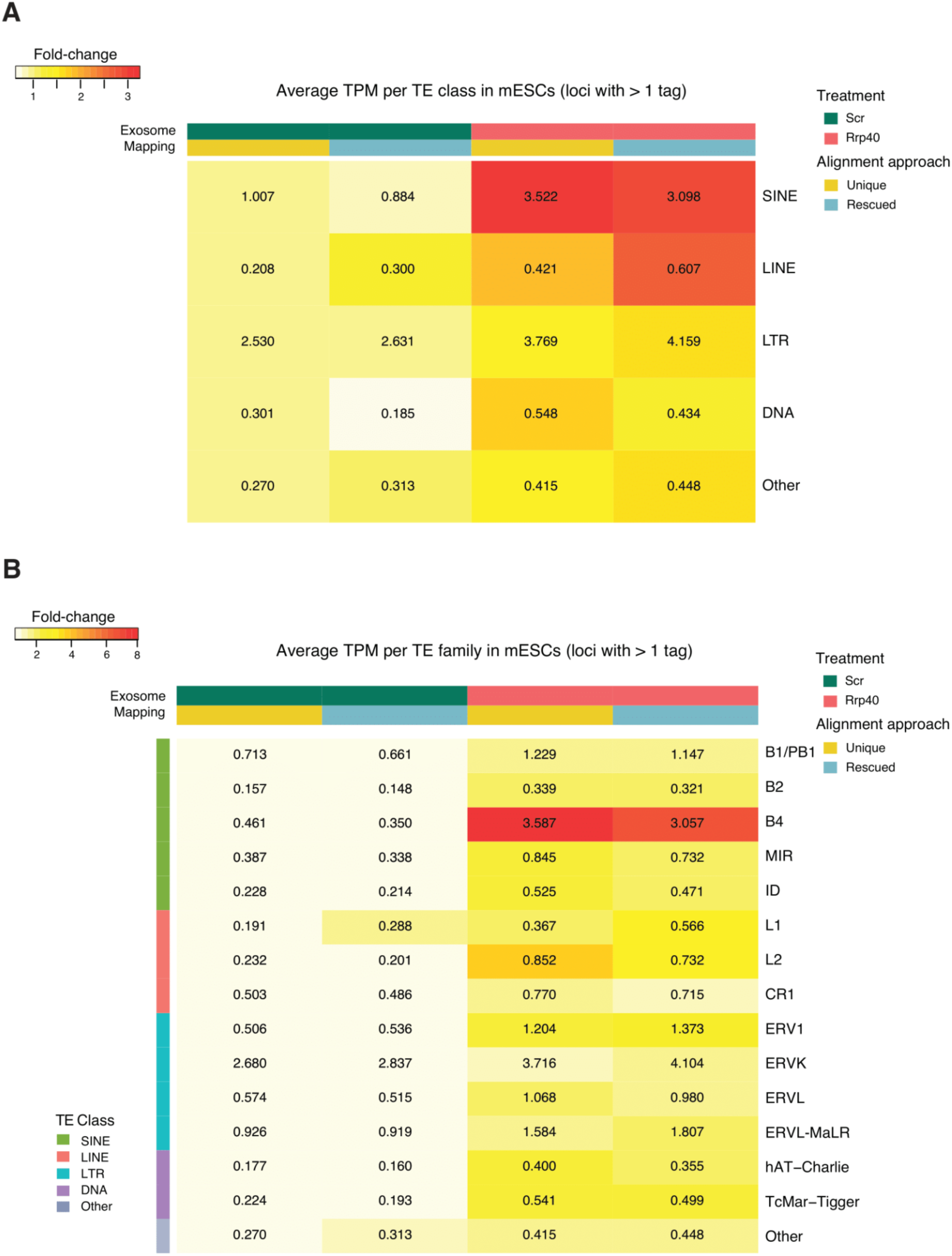
**A-B**: Heatmaps of average TPM-normalized expression values quantified at the CTSS level for major TE classes (**A**) and major TE families (**B**) in mESCs. Values for control (Scr) or exosome-depleted (Rrp40) CAGE libraries and using only uniquely mappable reads (Unique) or employing the MumRescueLite algorithm (Rescued) are shown per column. The values in the cells are calculated as TPM values divided by the union of TEs expressed in at least one of the columns/conditions, thus representing a comparable average TPM value. The color key represents the fold-change versus the value for uniquely mappable reads in control samples.

**Supplementary Figure 3.**
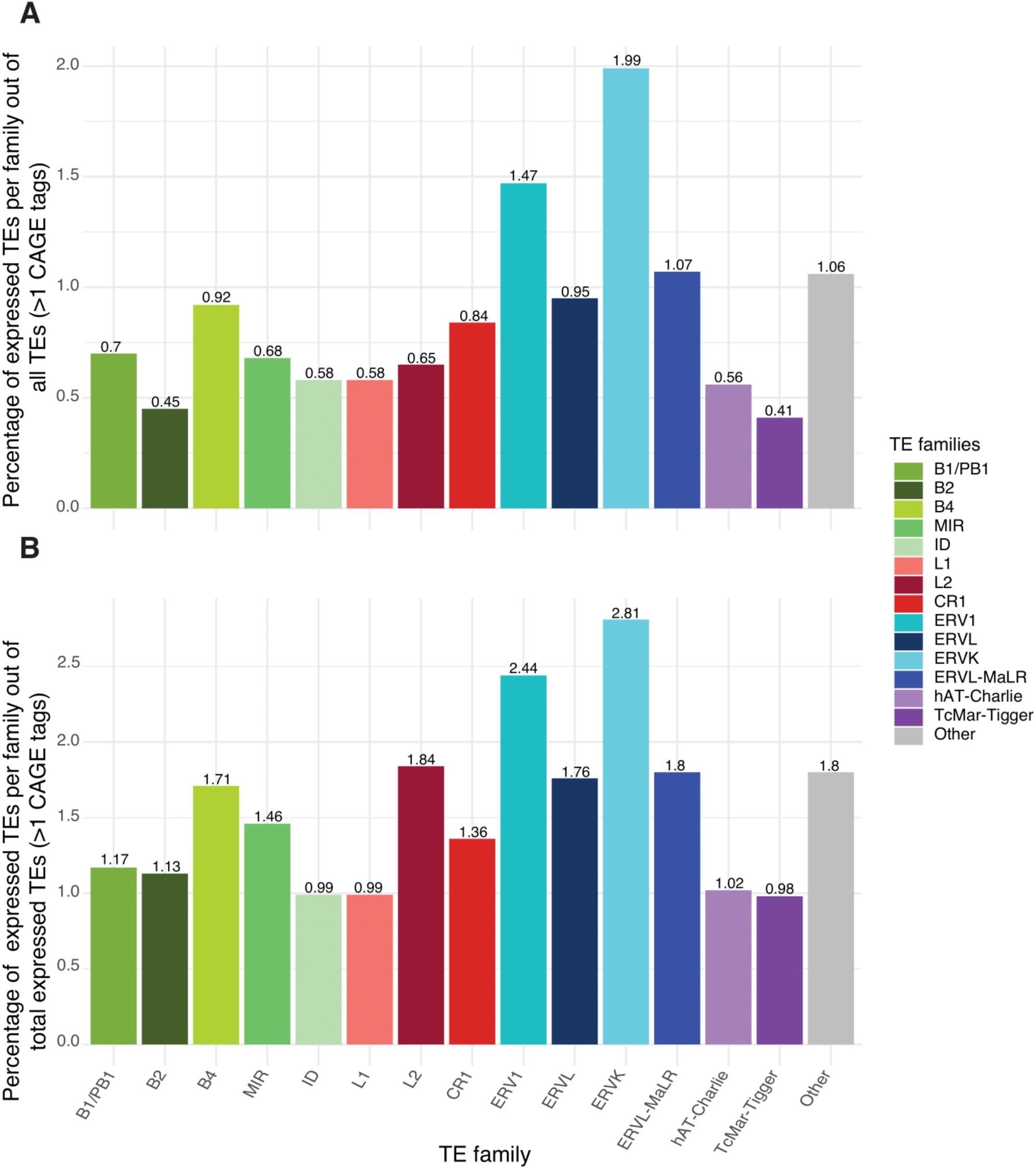
**A-B:** Percentage of transcribed TEs at the TE family level out of all TE insertions annotated in RepeatMasker (A) and out of all identified TEs with two or more CAGE tags (B).

**Supplementary Figure 4.**
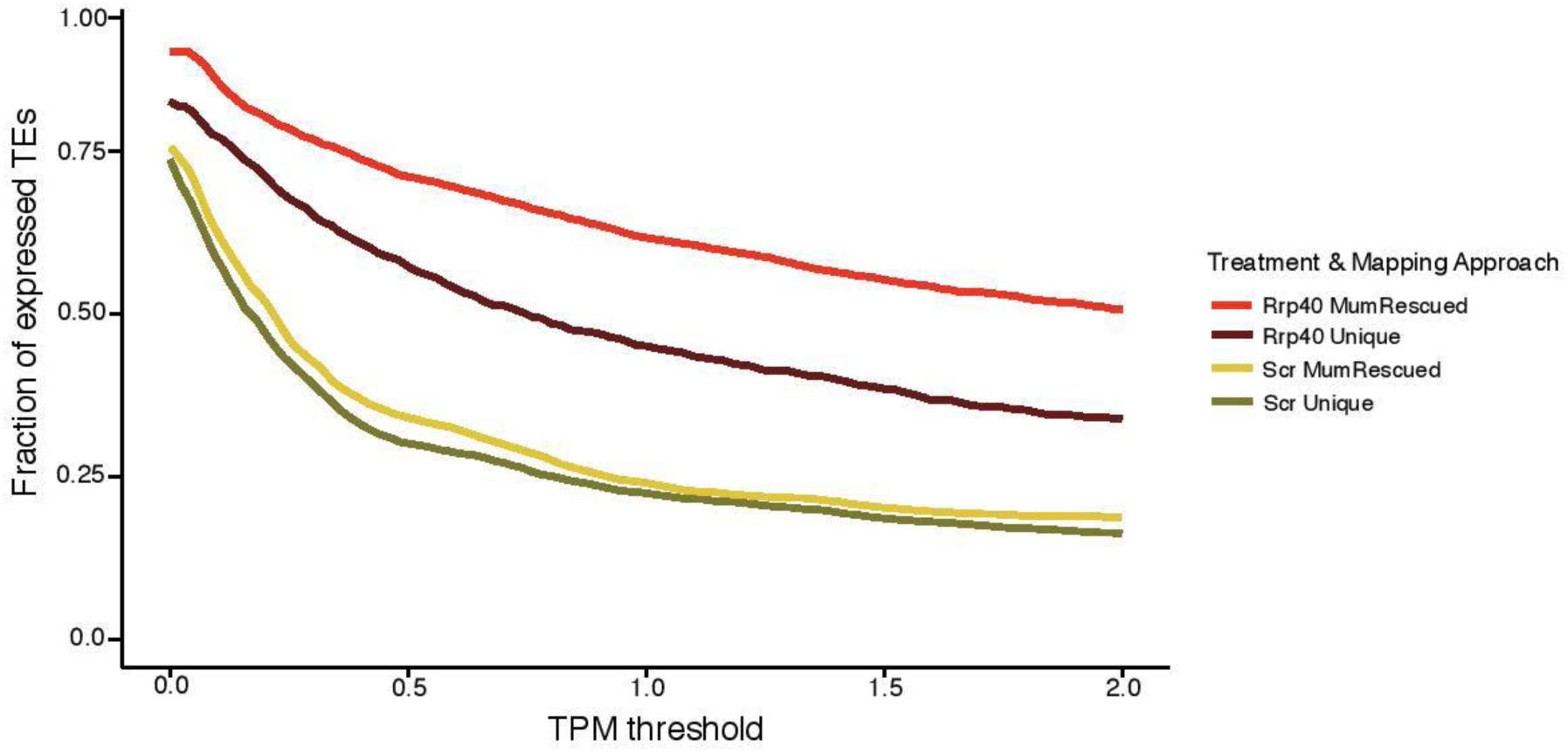
Fraction of expressed TE insertions versus TPM expression threshold (horizontal axis) over all expressed TEs in control (Scr) and exosome-depleted (Rrp40) libraries, counting only uniquely mappable (Unique) or, in addition, also rescued (MumRescueLite-processed) reads.

**Supplementary Figure 5.**
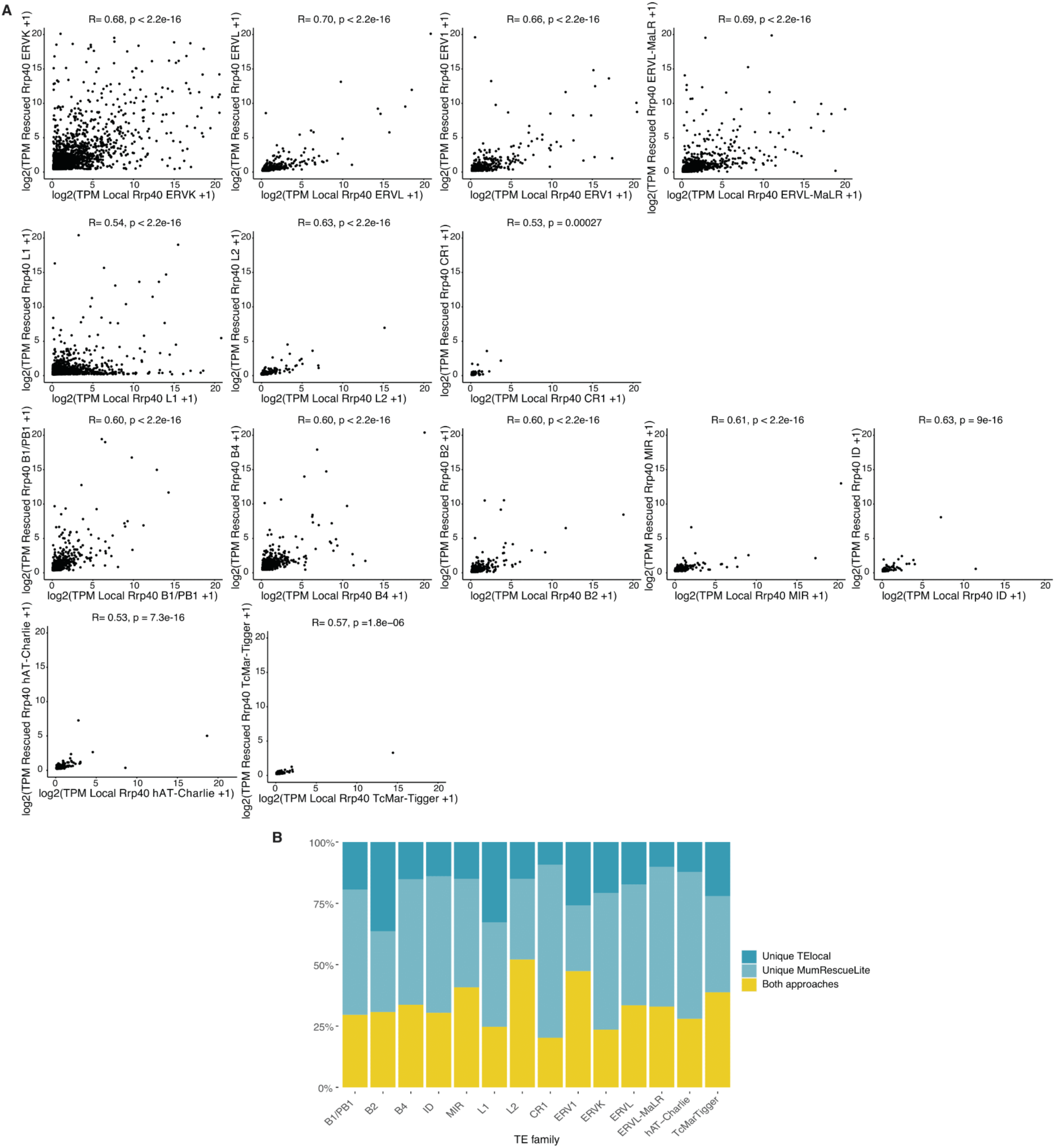
**A**: Quantified expression levels (log_2_-transformed TPM-normalized counts) by MumRescueLite (vertical axis) and TElocal (horizontal axis) from pooled CAGE libraries. Expressed TE insertions detected by both approaches for major TE families are shown. **B**: Proportions of expressed TEs identified by TElocal, MumRescueLite or both approaches for major TE families.

**Supplementary Figure 6.**
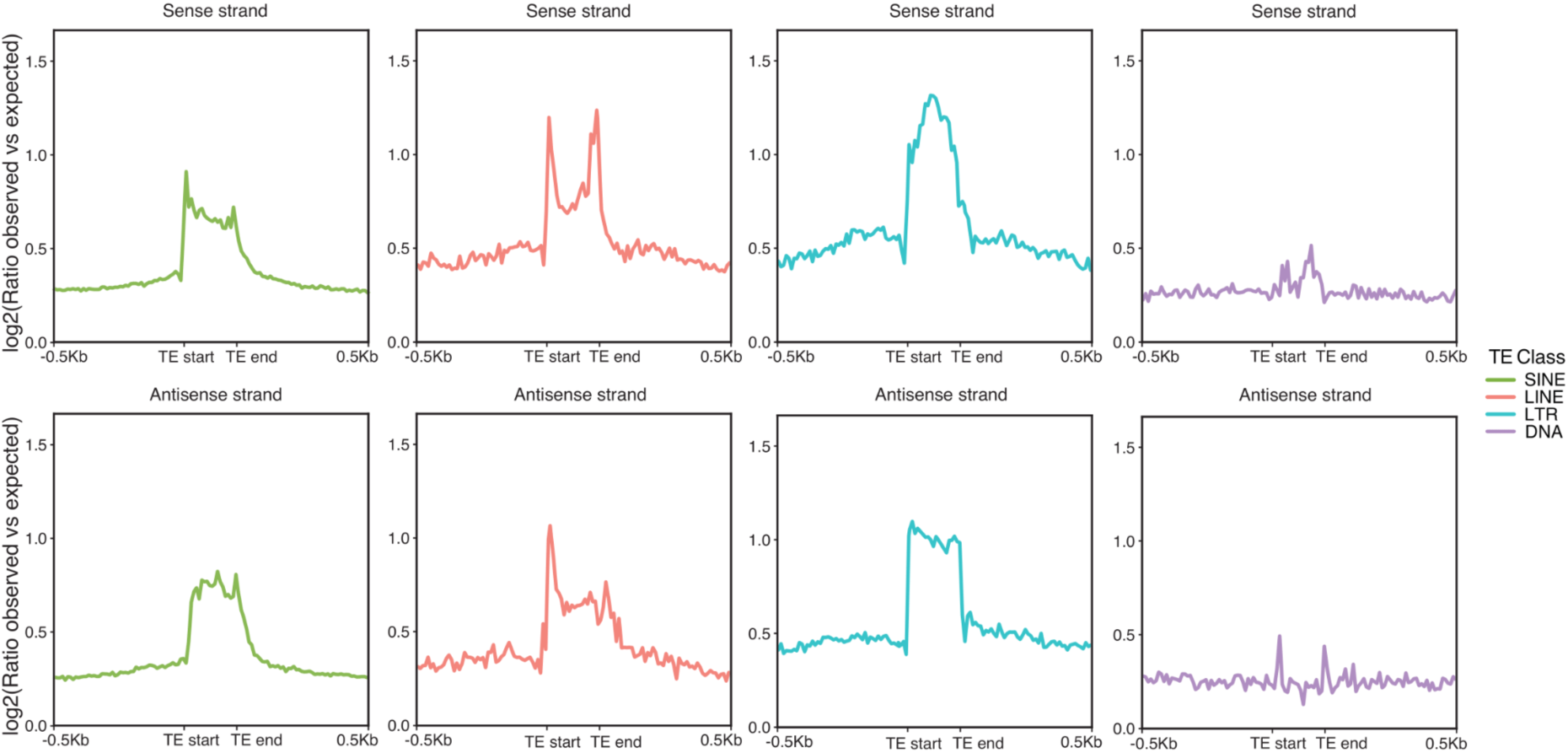
Log_2_ ratio of average observed versus expected (as determined by a synthetic CAGE uniqueness track, see Methods) distribution of CAGE-inferred TSS locations (vertical axis) +/- 500 bp upstream/downstream and across the body of major TE classes (horizontal axis). TSS locations are visualized separately for the sense (upper panel) and antisense (lower panel) strands.

**Supplementary Figure 7.**
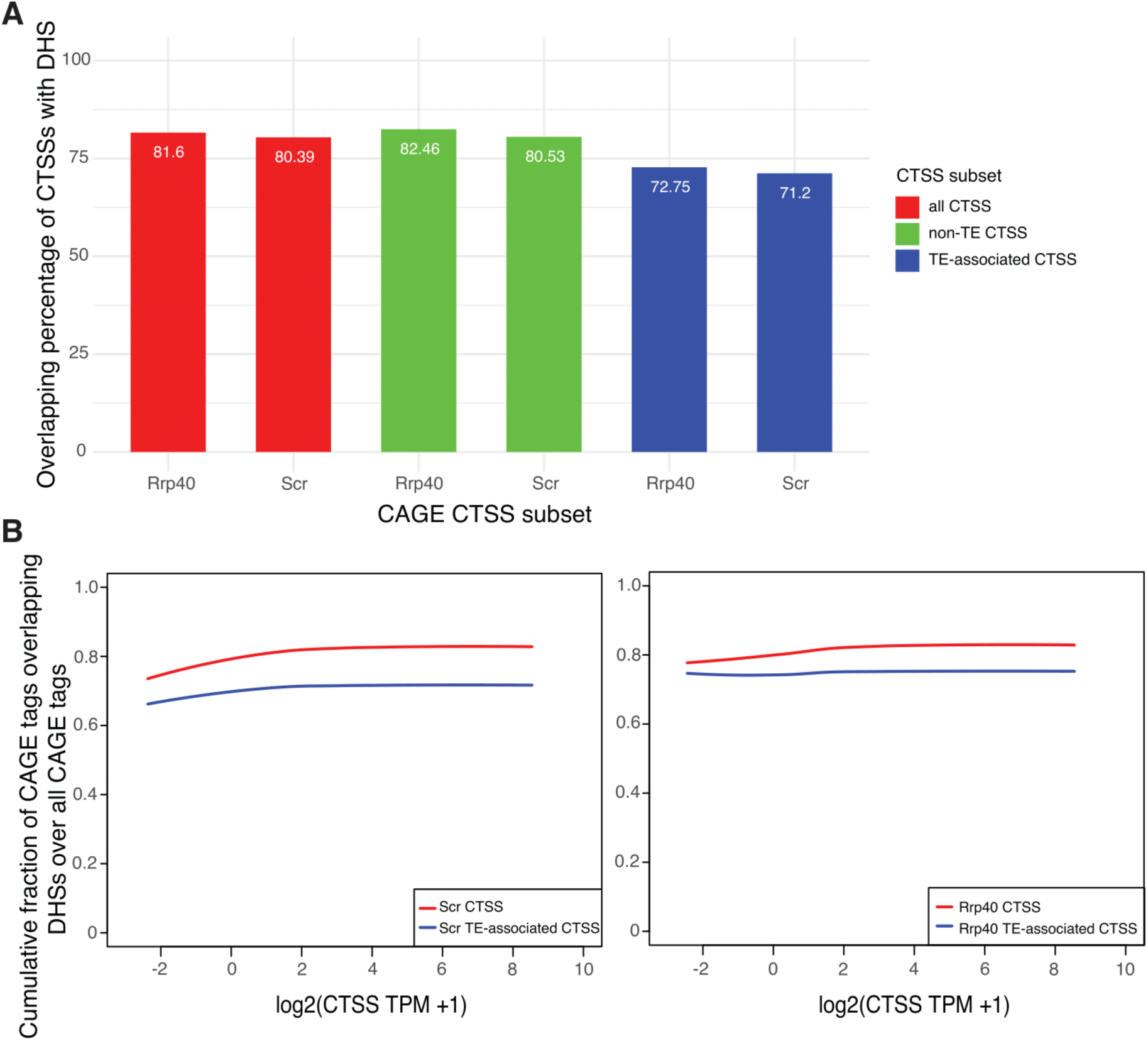
**A:** Percentage of CAGE CTSSs within +/- 250 bp of DHS midpoints out of total in each of the subsets: all called (red), all non-TE associated (green) and all TE-associated (blue) CAGE CTSS in control and exosome-depleted CAGE libraries, respectively. **B:** Cumulative fraction of CAGE tags within +/- 250 bp of DHS midpoints out of total CAGE ctss in control (Scr, left panel) and exosome-depleted (Rrp40, right panel) CAGE libraries versus CTSS expression level (log_2_-transformed TPM-normalized counts). The cumulative fraction for all CAGE CTSS (red) and TE-associated CTSSs (blue) is shown.

**Supplementary Figure 8.**
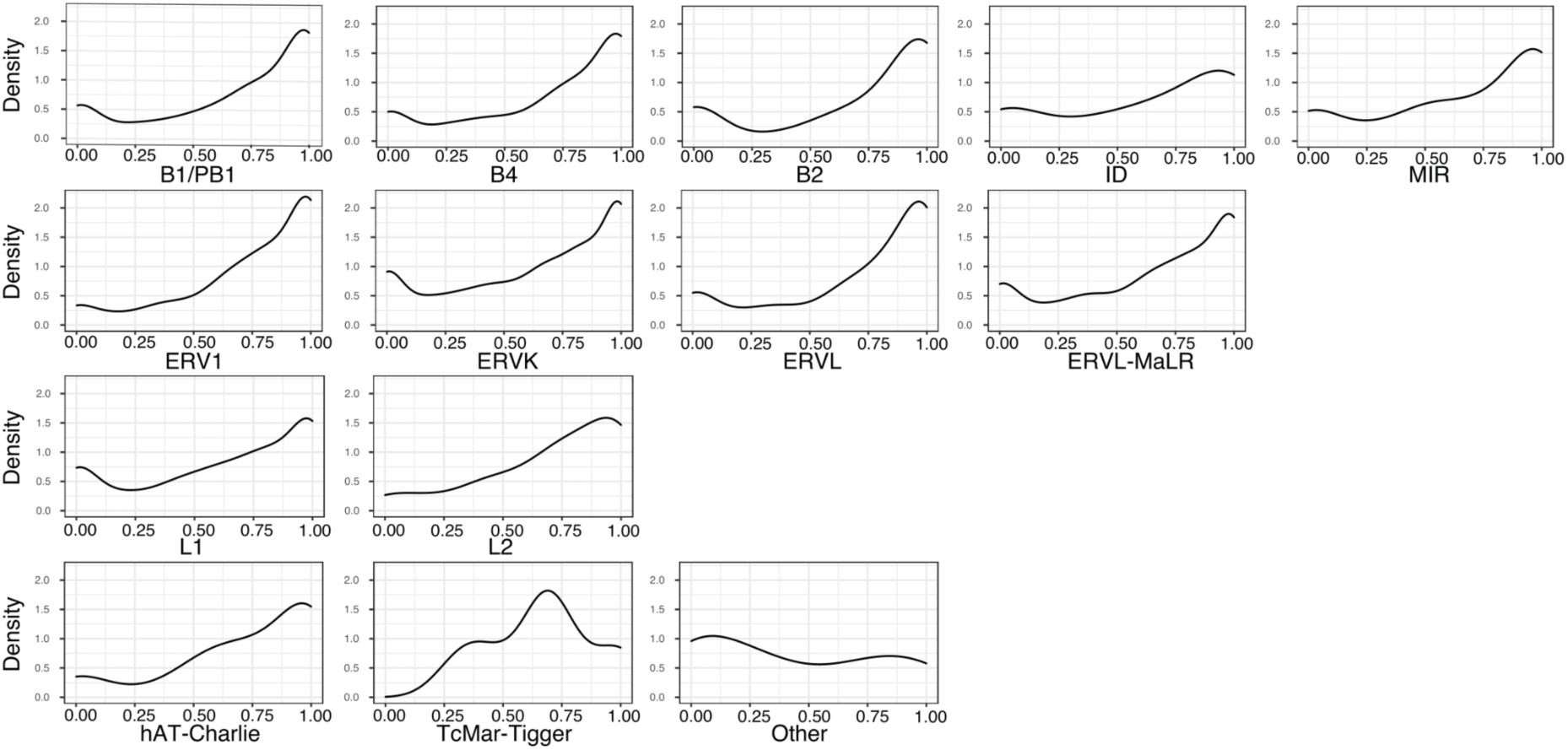
Densities of exosome sensitivity score, measuring the relative amount of exosome degraded RNAs, for transcripts associated with major TE families based on DHS-associated strand-specific expression levels in control and exosome-depleted CAGE libraries. The exosome sensitivity score ranges from 0 (fully captured by control CAGE) to 1 (only observed upon exosome depletion).

**Supplementary Figure 9.**
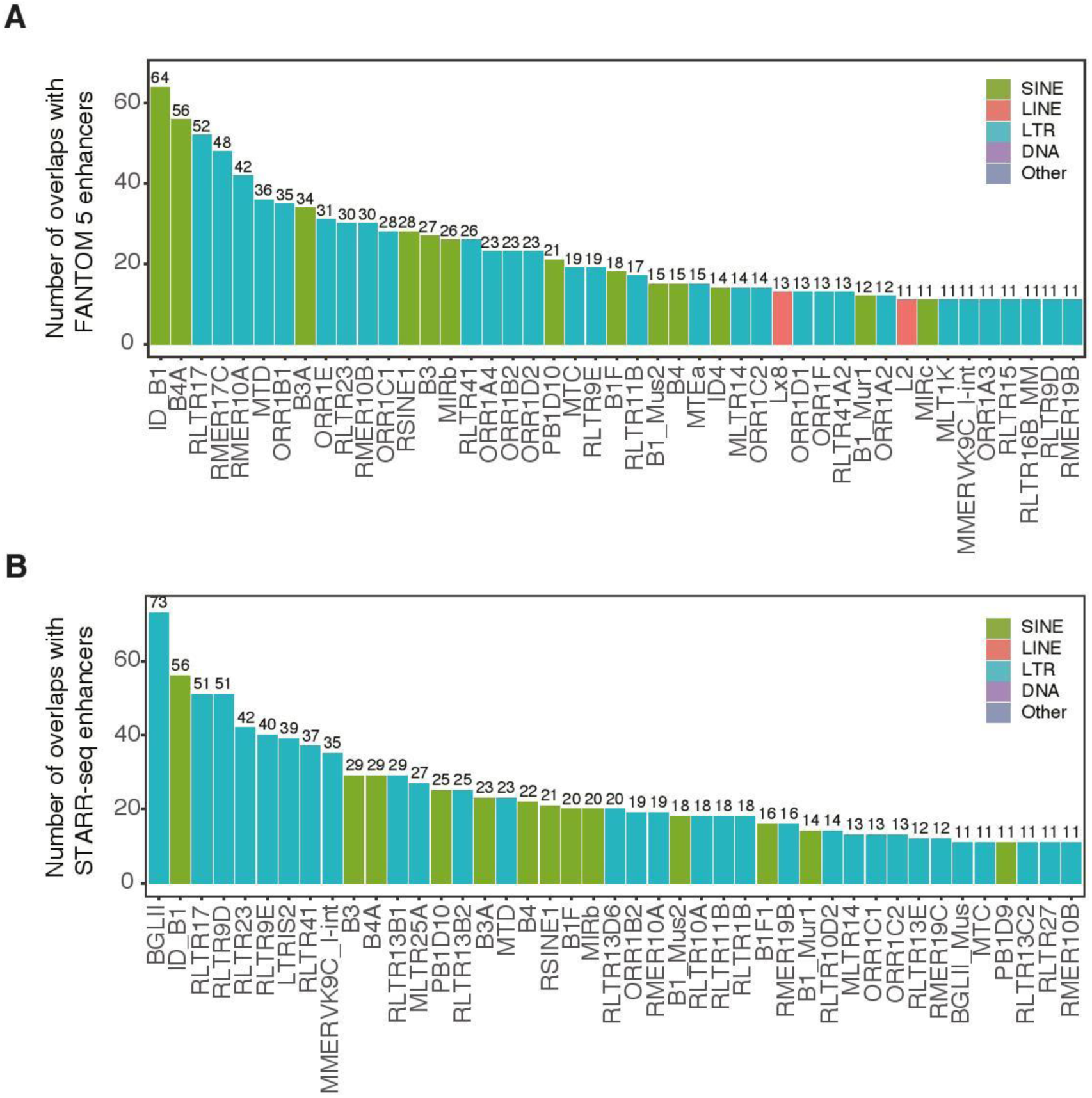
The number of transcribed TEs overlapping FANTOM5 mouse enhancers (A) and STARR-seq mESC enhancers (B) at the TE subfamily level.

**Supplementary Figure 10:**
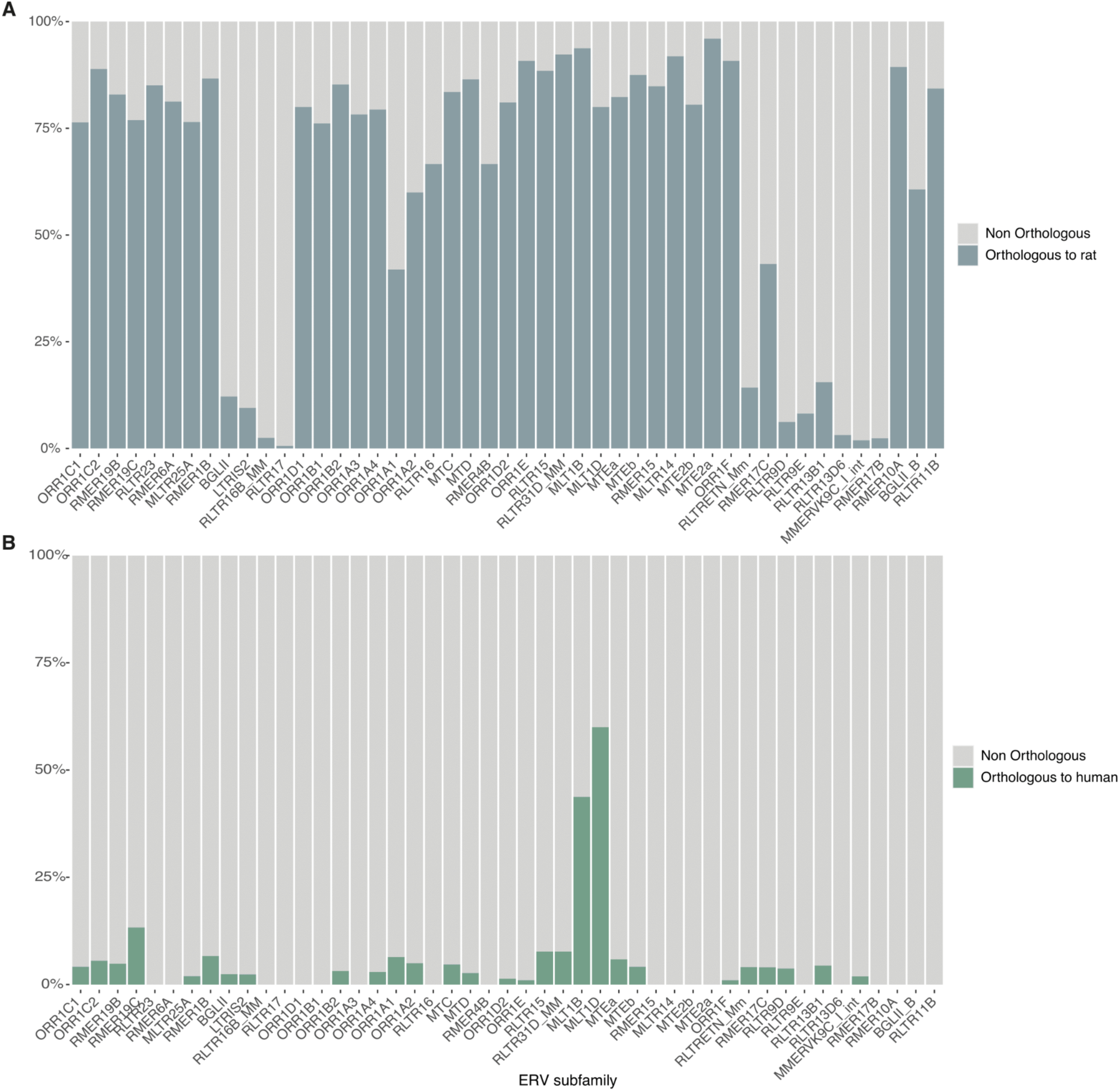
Proportion of mouse TE-associated expressed DHSs with orthologous regions in the rat (A) and the human (B) genomes split by ERV subfamily.

**Supplementary Figure 11:**
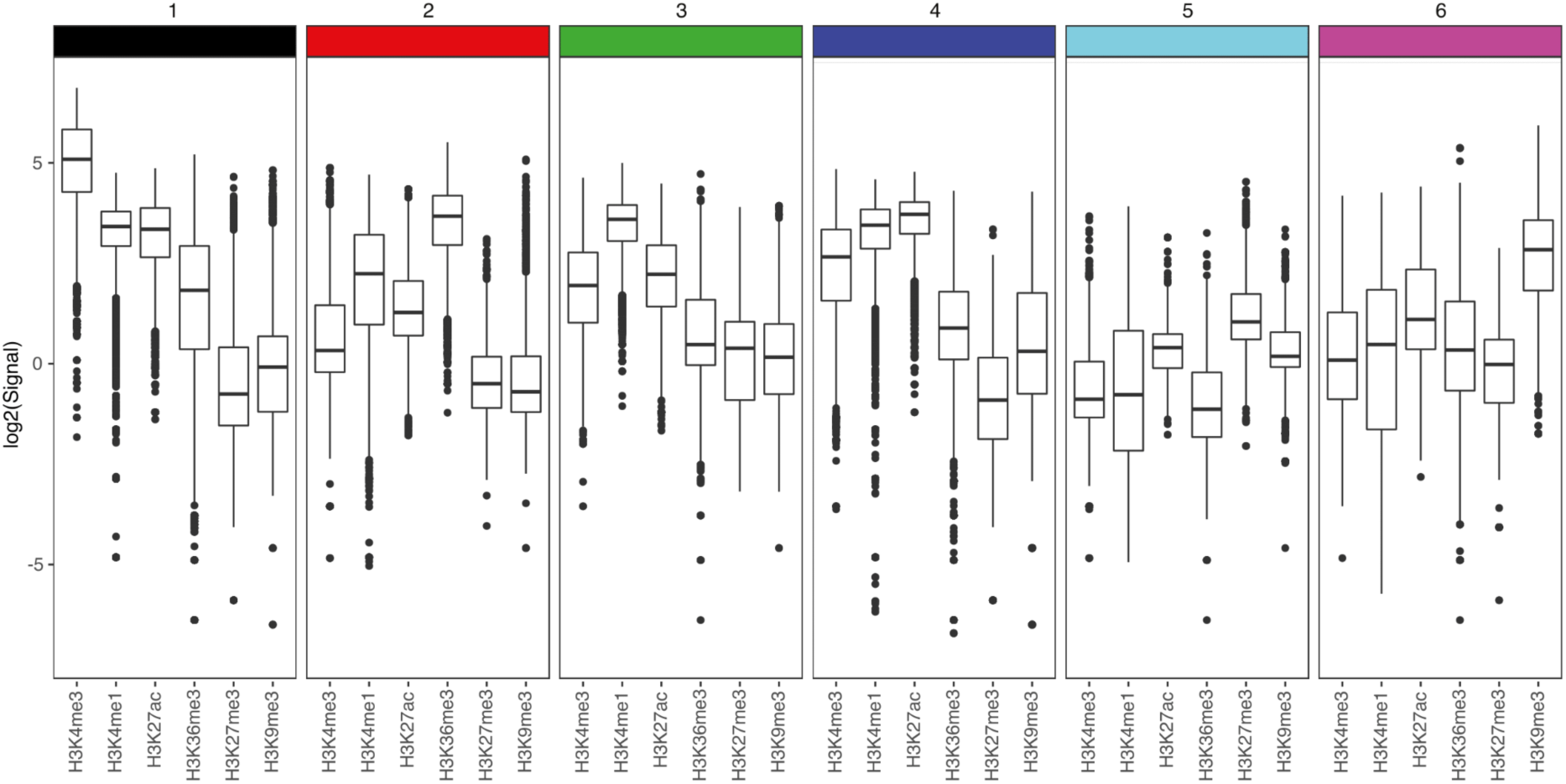
Box-and-whisker plots of log_2_-transformed histone modification (ChiP-seq) signal around summits of TE-associated clusters of CAGE-inferred TSSs for groups identified by hierarchical clustering, as shown in Figure 4A. Central band: median; boundaries: first and third quartiles; whiskers: +/- 1.5 IQR.

**Supplementary Figure 12:**
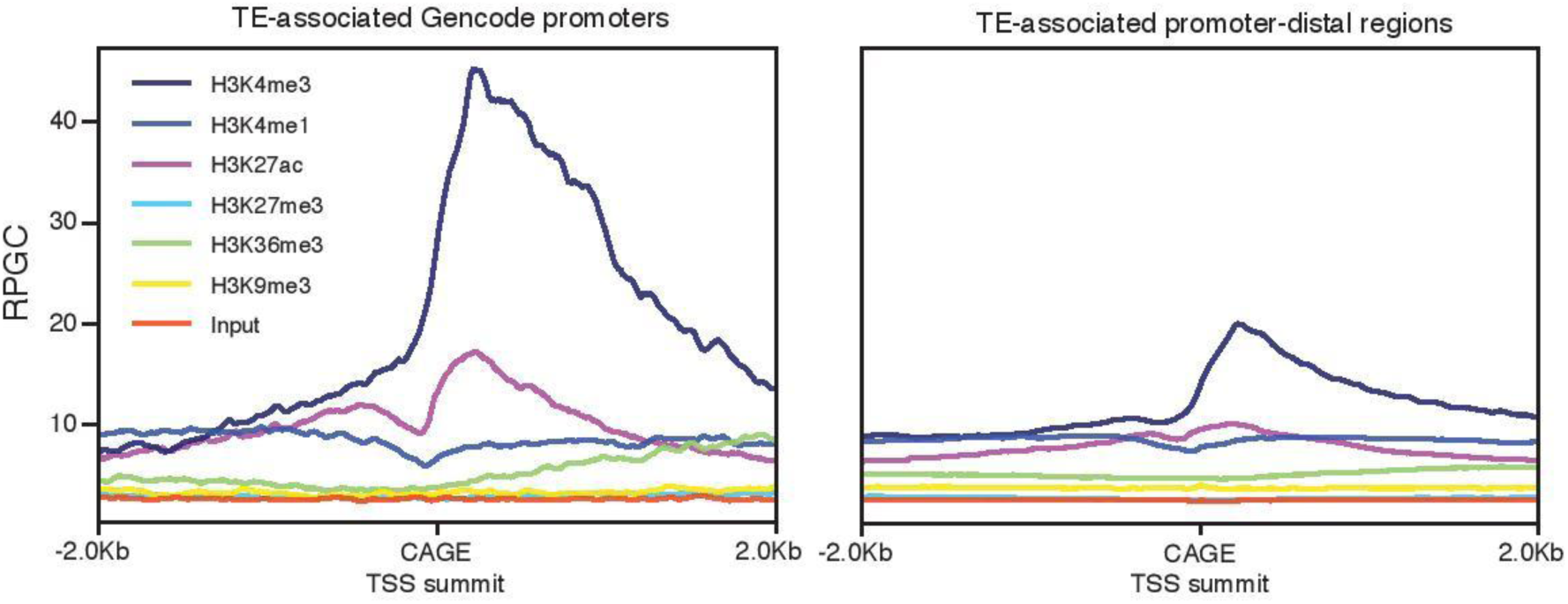
Average ChIP-seq signal for histone modifications (H3K4me3, H3K4me1, H3K27ac, H3K27me3, H3K36me3, H3K9me3) and input +/- 2 kb around the summits of TE-associated CAGE-inferred TSS clusters at GENCODE promoters (left) and at gene-distal locations (right). Signals shown are reads per genomic context (RPGC), normalized for sequencing depth to 1x genome coverage.

**Supplementary Figure 13.**
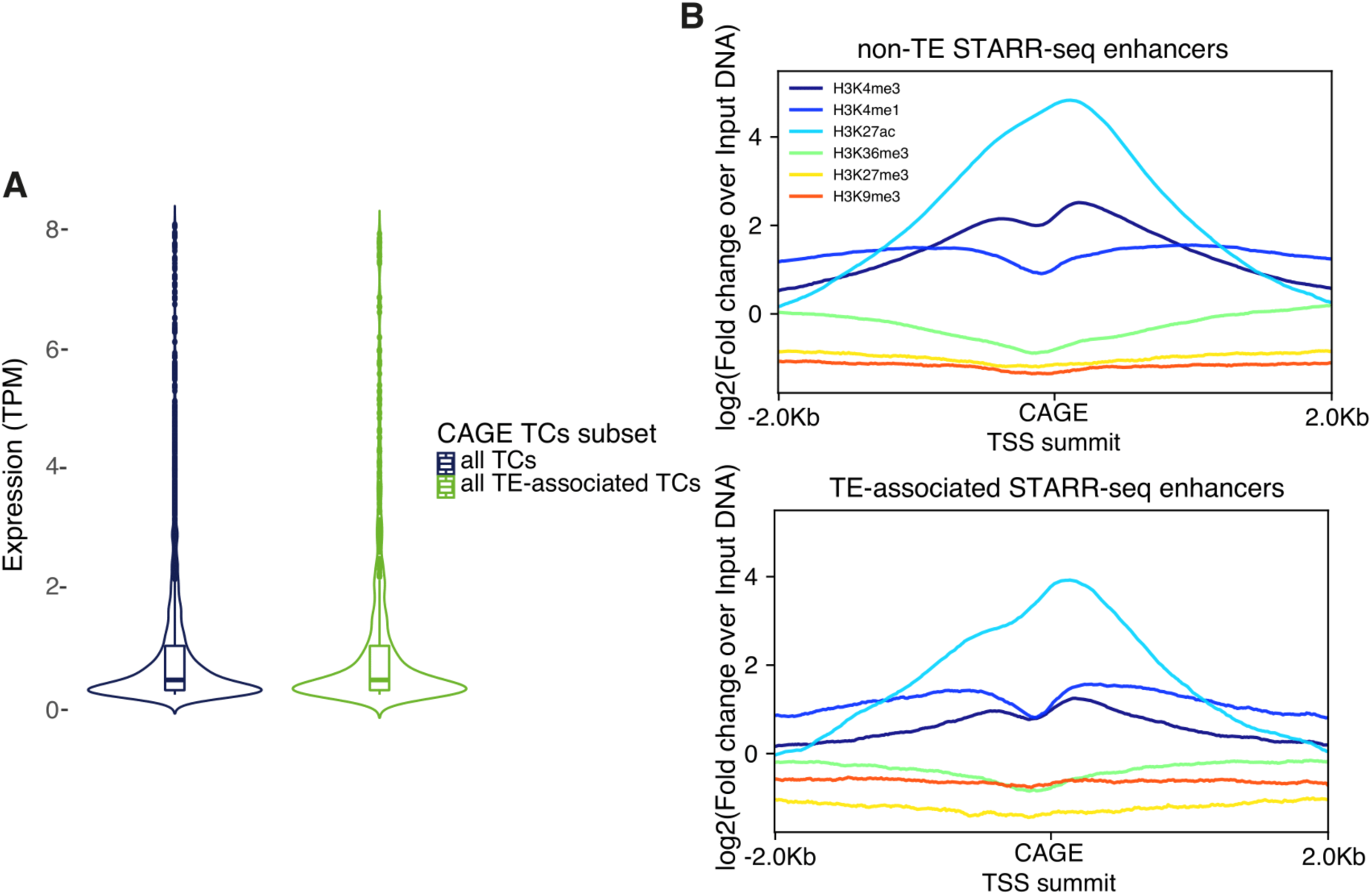
Comparison of TPM-normalized expression (A) and normalized ChiP-seq signal for histone modifications (fold change over input DNA, B) for TE- and non-TE-associated CAGE-inferred TSS clusters (TCs) overlapping DHSs as well as STARR-seq enhancer regions.

**Supplementary Figure 14.**
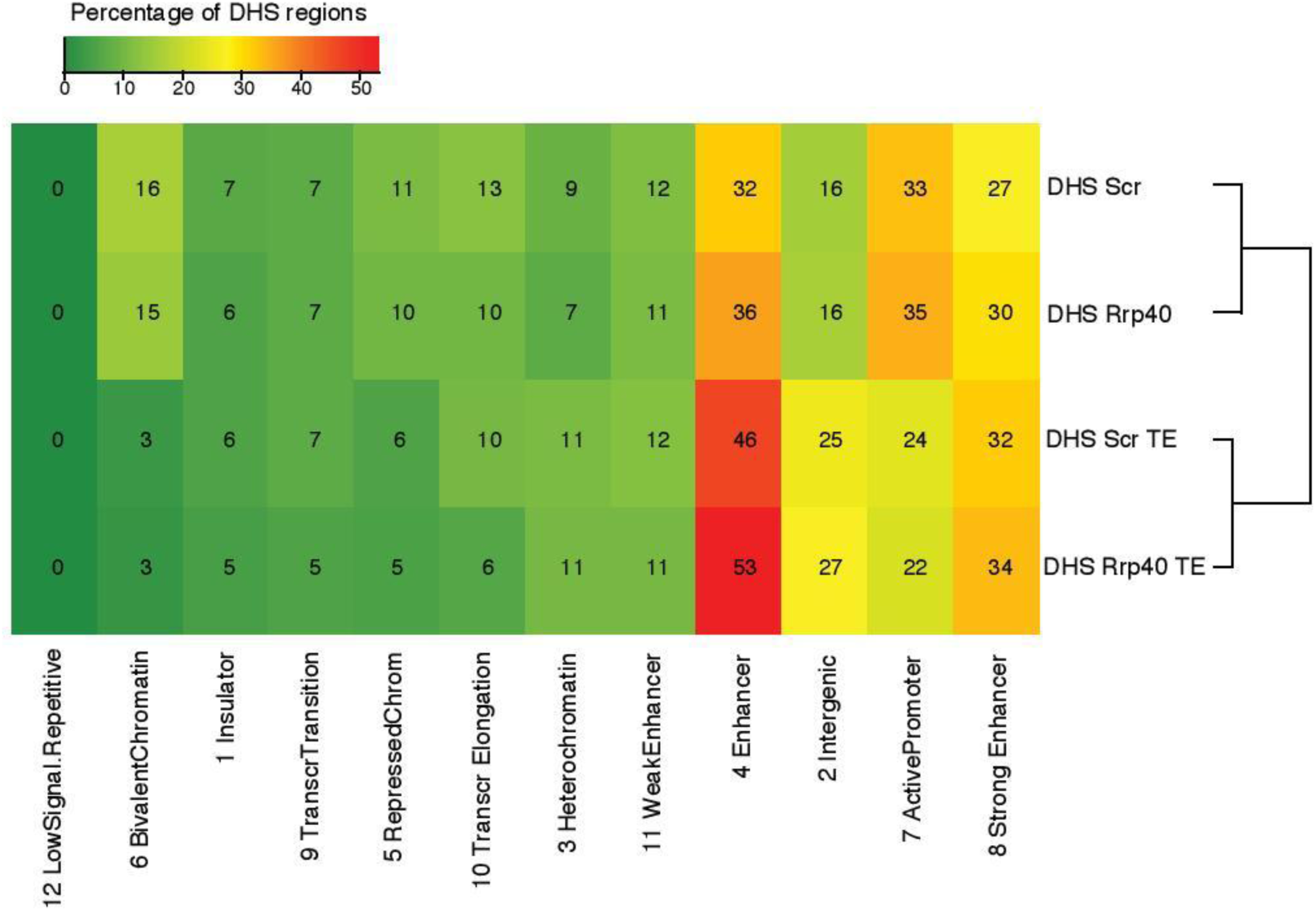
12-state ChromHMM model revealing the enrichment of specific combinations of histone modifications and TF marks at expressed DHS-associated genomic regions. The color key (from green (low) to red (high)) and the values in the heatmap cells show the percentage of DHSs that overlap each different ChromHMM state (horizontal axis) that are either expressed in control (Scr) and exosome-depleted (Rrp40) CAGE libraries, split by considering either all expressed DHSs or those that also overlap transcribed TEs.

**Supplementary Figure 15.**
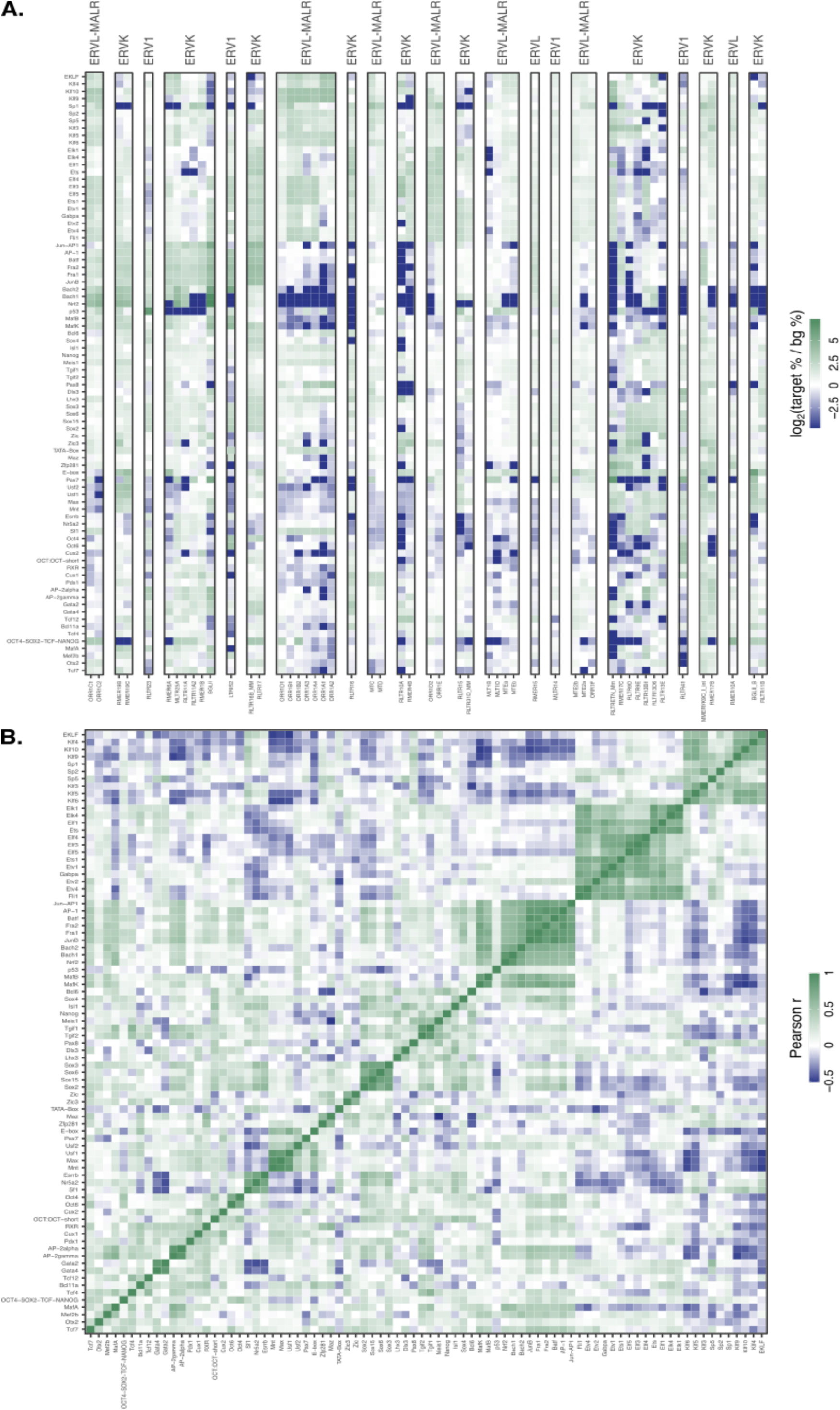
**A:** Motif enrichments for TFs (columns) in TE-associated transcribed DHSs for all ERV subfamilies (rows) versus a background of genomic regions +/- 200 bp around the summits of all CAGE-inferred TSS clusters. White cells indicate either no enrichment or non-reported values. **B:** Pearson’s correlation coefficients for pairwise TF motif co-occurence across all ERV subfamilies.

**Supplementary Figure 16.**
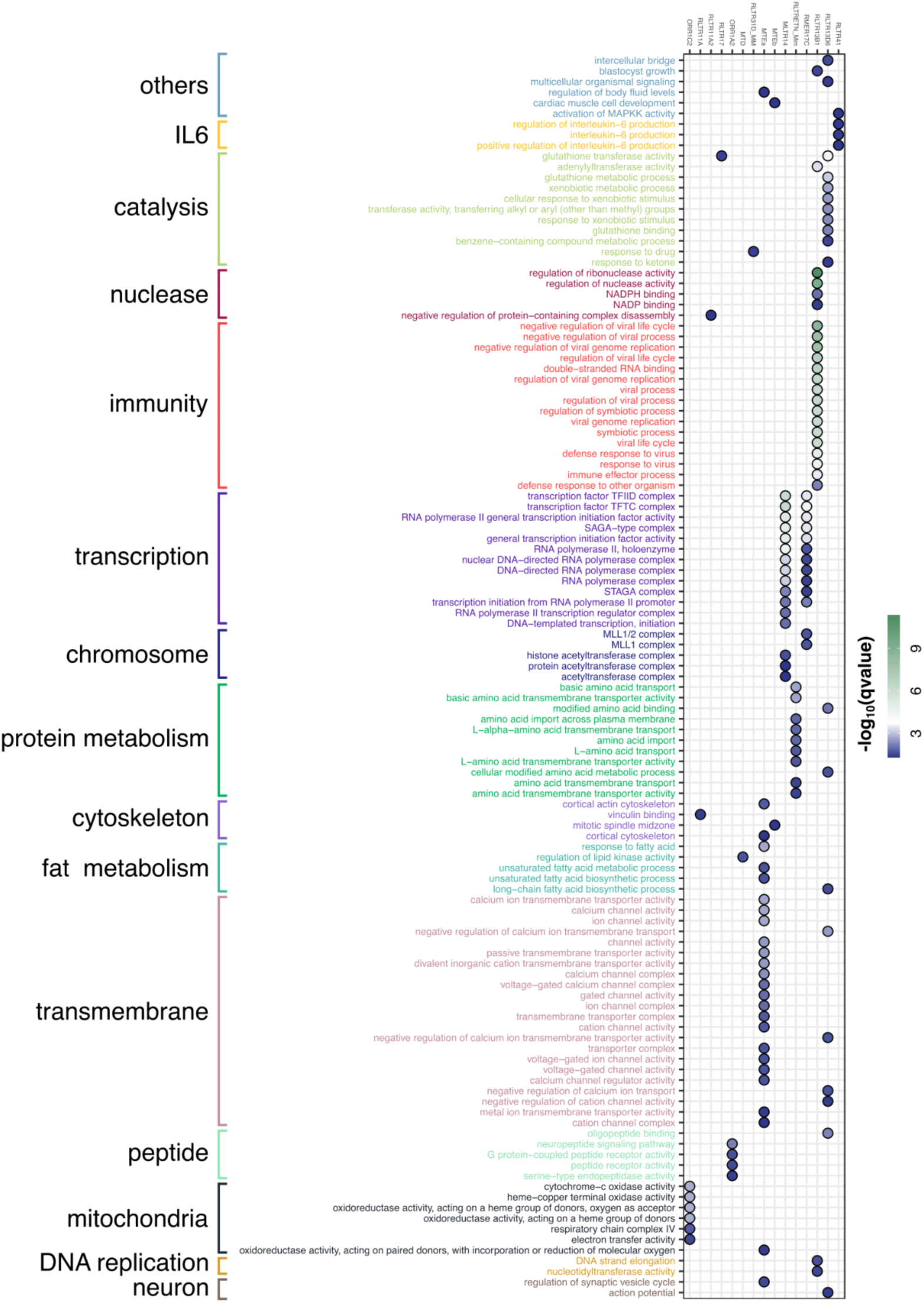
Enriched GO terms based on putative target genes (ABC) of gene distal ERVs for individual ERV subfamilies (foreground, rows) versus all ABC-predicted target genes (background). All enriched GO terms are manually curated into processes or functions and colored accordingly. Points are colored according to the transformed q-values indicating the significance of the GO-term enrichment.

